# NPC1 deficiency engages a lysosome–genome–immune program linked to neurodegeneration and cellular aging signatures

**DOI:** 10.64898/2026.07.12.738054

**Authors:** Morteza Abyadeh, Masoumeh Zarei Kheirabadi, Pei-Chi Hou, Vedat Sari, Kyubin Lee, Rohil Hameed, Bushra Mehkri, Elizabeth Malaugh, Jason Newton, Tomasz Kordula, Yuh-Hwa Wang, Alaattin Kaya

## Abstract

Lysosomal dysfunction is a prominent feature of neurodegeneration and aging, yet how primary defects in lysosomal trafficking are converted into progressive cellular decline remains poorly understood. Niemann Pick disease type C (NPC), caused by impaired *NPC1* dependent cholesterol export, provides a genetically defined model to address this question. Here, we show that NPC1 deficiency activates a lysosome, genome, immune axis linking cholesterol trafficking failure to neurodegeneration and hallmarks of cellular aging. In Npc1 mutant mice, *NPC1* loss triggered DNA damage, neuroinflammation, microglial and astrocytic activation, Purkinje neuron degeneration, and motor dysfunction. Consistently, NPC patient-derived fibroblasts exhibited mitochondrial abnormalities and widespread DNA double-strand breaks. Genome-wide DNA break mapping and transcriptomic analyses revealed extensive genomic instability at regulatory regions, including enrichment of DNA breaks at transcription start sites and G quadruplex associated loci, accompanied by widespread transcriptional reprogramming, activation of innate immune pathways, disruption of fibroblast identity, and induction of cellular aging signatures. We further identify Fingolimod, an FDA approved sphingosine - 1 phosphate receptor modulator, as a potent modifier of this disease network. Fingolimod improved lysosomal cholesterol trafficking, increased *LAMP1* abundance, attenuated *STING* associated inflammatory signaling, normalized mitochondrial function, reduced neuroinflammatory and neurodegenerative phenotypes in Npc1 mutant mice, and broadly shifted disease-associated transcriptional programs toward a healthier state. Extending these findings beyond NPC, Fingolimod improved age-associated phenotypes in *C. elegans* and prolonged lifespan in aged male mice. Together, these findings identify genome instability and chronic innate immune activation as major downstream consequences of lysosomal cholesterol trafficking failure and establish Fingolimod as a clinically actionable modulator of lysosomal dysfunction, neurodegeneration, and aging-related decline.

## INTRODUCTION

Lysosomes are essential organelles that maintain cellular homeostasis through the degradation and recycling of proteins, lipids, nucleic acids, and damaged organelles ^1^. Impairment of lysosomal function results in the accumulation of toxic substrates, defective organelle turnover, metabolic imbalance, and cellular stress ^2^. Genetic defects affecting lysosomal proteins give rise to lysosomal storage diseases (LSDs), a heterogeneous group of inherited disorders characterized by progressive substrate accumulation and multi-organ pathology, including involvement of the nervous system ^3^. Beyond classical LSDs, lysosomal dysfunction is now recognized as a broader hallmark of aging and neurodegeneration ^1^. Recent advances in transcriptomics, single-cell omics, CRISPR-based modeling, and disease-relevant induced pluripotent stem cell systems have further highlighted the lysosome as a central hub integrating cellular metabolism, neuroinflammation, and stress-response pathways in aging and disease ^4^. Intriguingly, transcriptomic aging-clock analyses have revealed that Niemann–Pick disease type C (NPC), a monogenic lysosomal storage disorder, exhibits features of accelerated brain aging, suggesting that lysosomal failure may actively drive aging-associated molecular programs and neurodegenerative processes ^5^. These findings position lysosomal dysfunction not only as a cause of rare inherited disorders, but also making NPC a genetically defined model for studying how lysosomal dysfunction contributes to aging and neurodegeneration.

NPC is a rare, autosomal recessive lysosomal storage disorder caused by mutations in *Npc1* (∼95%) or *Npc2* (∼5%), genes that encode proteins essential for the transport of unesterified cholesterol from the late endosomal/lysosomal (LE/LY) compartment ^[6]^. Loss of NPC1 or NPC2 function results in the pathological accumulation of cholesterol and other lipids, including glycolipids and fatty acids, within LE/LY organelles ^[7^,^8]^. This primary defect in lipid trafficking results in progressive multisystem disease, with prominent neurological manifestations including supranuclear gaze palsy, motor impairment, seizures, and cognitive decline ^[9]^. Although NPC has a broad age of onset ranging from infancy to adulthood, neurological onset most commonly occurs in childhood and is strongly associated with disease severity and progression, frequently culminating in premature mortality ^[9^,^10]^.

Accumulating evidence indicates that impaired cholesterol trafficking, lysosomal dysfunction, and chronic neuroinflammation are central drivers of NPC pathogenesis. Consistent with this view, therapeutic interventions aimed at restoring lipid homeostasis or suppressing neuroinflammatory signaling significantly ameliorate disease progression in NPC1-deficient mouse models ^[11^,^12]^. In late 2024, two therapies, miplyffa and aqneursa, received FDA approval for the treatment of neurological manifestations associated with NPC. While these advances represent a major milestone, their mechanisms of action remain incompletely understood, and both drugs are costly and associated with adverse effects, including infections, gastrointestinal symptoms, and weight loss ^[13^,^14]^. These limitations highlight the need for safer, more accessible therapeutic strategies that target core disease mechanisms.

Beyond lipid storage, NPC pathology is characterized by widespread cellular dysfunction, including neuroinflammation, selective neuronal vulnerability, axonal degeneration, and hypomyelination ^6^. These features are shared with multiple neurodegenerative disorders, underscoring lysosomal failure as a convergent pathogenic mechanism rather than a disease-specific anomaly ^4^. However, unlike more common neurodegenerative diseases, NPC represents a genetically defined model in which lysosomal dysfunction is the primary initiating event, thereby providing a unique opportunity to dissect downstream pathological cascades that have remained largely elusive.

Given two key pathological hallmarks of the disease including inflammation and dysfunctional lipid metabolism, we sought to investigate the therapeutic potential of Fingolimod, an FDA-approved drug with established anti-inflammatory properties and emerging roles in lipid metabolism, as a repurposed treatment for NPC. Fingolimod was developed from myriocin, a bioactive compound isolated from *Isaria sinclairii*, a fungus used in traditional Chinese medicine ^7^. It is FDA approved for multiple sclerosis, functions as a sphingosine-1-phosphate (S1P) receptor modulator, thereby reducing neuroinflammation and modulating immune responses key pathological features shared by multiple sclerosis and NPC ^6,8,9^. Fingolimod has also been shown to partially correct cholesterol trafficking defects in human *Npc1* mutant fibroblasts by reducing cholesterol and sphingolipid accumulation and improving intracellular lipid trafficking, however, its *in vivo* efficacy and mechanistic basis remain elusive ^10^. In addition, a recent clinical study demonstrated that Fingolimod treatment in patients with Neuronal ceroid lipofuscinosis type 1 (CLN1) disease, another rare autosomal recessive lysosomal storage disorder, resulted in a significant reduction in neurofilament light chain (NfL) levels, a biomarker of neuronal injury and axonal degeneration, although the underlying mechanism of action remains poorly understood ^11^.

In this study, we investigated the effects of Fingolimod in NPC using *Npc1*-mutant mouse models and patient-derived fibroblasts, and extended our analyses to additional model systems, including *Caenorhabditis elegans* and aged mouse cohorts. Through integrated molecular, cellular, and transcriptomic analyses, we identified disease-associated alterations linking lysosomal dysfunction to neuroinflammation, mitochondrial abnormalities, genome instability, and cellular aging-related signatures that have not been previously described. We found that Fingolimod partially ameliorated lysosomal defects, as reflected by increased LAMP1 abundance and improved intracellular cholesterol trafficking, while attenuating inflammatory signaling and improving key neurodegenerative phenotypes in NPC models, despite having limited effects on DNA damage. Notably, Fingolimod was associated with improved lifespan and age-related phenotypes across the model systems examined. Collectively, these findings suggest that lysosomal dysfunction contributes to multiple aspects of NPC pathology and supports further investigation of Fingolimod as a repurposed therapeutic candidate for NPC and aging-associated disorders.

## RESULTS

### Fingolimod Treatment Resulted in Delayed Weight Loss, Improved Motor Neuron Function and Preserved Neuronal Structure in Male *Npc1*^mut^mice

Because of the rarity of the disease, human studies are largely limited to patient-derived fibroblasts, macrophages, and induced pluripotent stem cell–derived neuronal cultures or brain organoids. In contrast, mouse models of NPC with the most common human *NPC1* mutation (*Npc1*^mut^: *Npc1^tm(I1061T)Dso^*), have been shown to robustly recapitulate key aspects of the human disease phenotype and are widely used to investigate disease pathogenesis *in vivo* ^12,13^. Hallmark features observed in mouse models include progressive weight loss and motor dysfunction, driven by neurodegeneration and characteristic histopathological changes in the brain, particularly in the cerebellum and Purkinje neurons ^12,14^. Accordingly, in this study we employed an established NPC mouse model to evaluate the effects of Fingolimod on body weight, motor function, and cerebellar Purkinje neuron integrity across experimental groups.

Fingolimod was administrated by oral gavage for 3 months starting from 20 days of age through day 105 with a dose of 2mg/kg/every other day and the weight was assessed every week. Both male and female *Npc1*^mut^ mice displayed weight loss at 8 weeks of age. Although, the effect is mild, Fingolimod treatment significantly delayed weight loss in males but not in females (**Fig. 1A, B**), Weight loss in NPC mice has been attributed to a combination of progressive neurodegeneration, impaired motor coordination, dysphagia, and increased metabolic burden associated with systemic inflammation and organ dysfunction ^15,16^. However, existing data on the impact of Fingolimod on body weight remain controversial, with studies reporting both weight loss and weight gain depending on the disease model. For instance, Fingolimod-induced weight loss has been observed in high-fat diet-induced obese mice ^17^, whereas weight gain has been reported in an acetic acid-induced ulcerative colitis rat model ^18^. These findings suggest that Fingolimod may modulate weight in a disease-dependent manner, potentially contributing to overall disease improvement. To further evaluate the therapeutic effects of Fingolimod on neurological function, motor performance was assessed using the balance beam and rotarod tests at postnatal day 90. In comparison to the age- and sex-matched non treated controls, both male and female *Npc1*^mut^ mice displayed significantly impaired motor function (**Fig. 1C-F**). When treated with Fingolimod, significant improvement for motor function was observed only for male *Npc1*^mut^ mice in comparison to the untreated male *Npc1*^mut^ controls (**Fig. 1C, D)**. Although, we also observed positive trend of improvement in both rotarod and balance beam test for female group, the effect was statistically insignificant (**Fig. 1E, F)**. It is important to note that, due to the small sample size (three mice per group), the behavioral test results should be interpreted with caution, as even a single mouse difference had a substantial impact on statistical significance.

**Figure 1.**
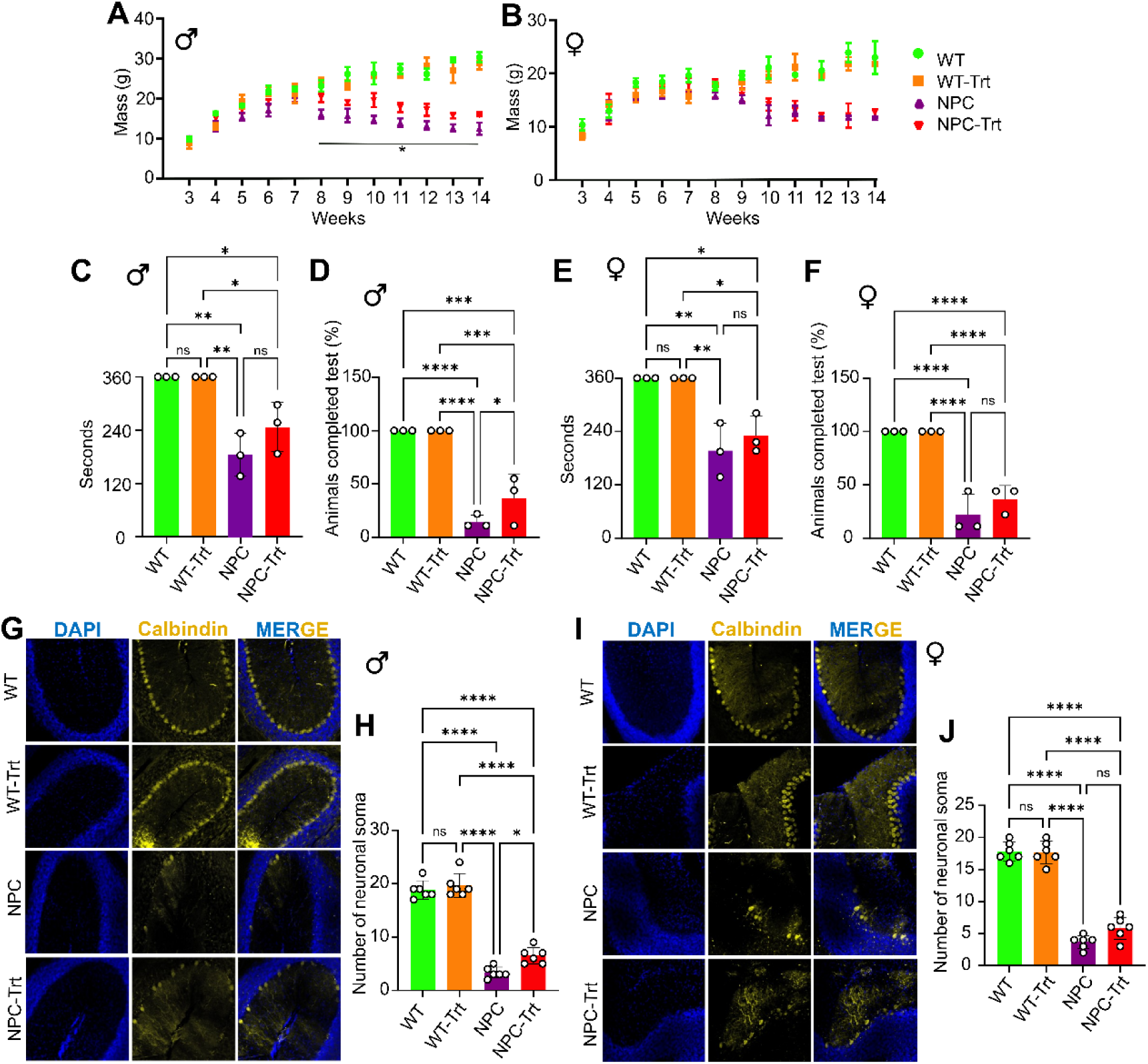
Fingolimod improves weight maintenance, motor performance, and Purkinje neuron integrity in male Npc1mut mice. (**A, B**) Body weight trajectories of male and female mice from week 3 to week 15 (day 105). Fingolimod treatment significantly attenuated weight loss in male Npc1mut mice between weeks 8 and 14 (**A**), while no significant effect was observed in females (**B**). (**C-F**) Motor performance assessed by rotarod and balance beam tests in male and female mice, respectively. Fingolimod-treated male Npc1mut mice exhibited improved motor function compared to untreated Npc1mut controls (**C, D**), with no significant differences observed in females. (**G-J**) Representative confocal images of cerebellar Purkinje neurons (Calbindin, yellow) and nuclei (DAPI, blue) i male (**G**) and female (**I**) mice across four groups: wild type (WT), WT treated with fingolimod (WT-Trt), Npc1mut (NPC), and Npc1mut treated with fingolimod (NPC-Trt). Quantification of Purkinje neuron numbers shows a significant reduction in Npc1mut mice, which is partially rescued by fingolimod treatment in males (**H**), but not in females (**J**). Images were acquired at 20× magnification. Data are presented as mean ± SEM. *p < 0.05, **p < 0.01, ***p < 0.001.

Progressive neuronal loss in the cerebellum is one of the main pathological events in NPC disease, where particularly cerebellar Purkinje cells (PCs) are affected ^19^. Purkinje cell loss in the cerebellum impairs motor coordination, balance, and motor learning by disrupting inhibitory output to deep cerebellar nuclei, leading to cerebellar ataxia. Loss of these cells causes motor dysfunction, such as gait abnormalities and tremors, and hinders the acquisition of new motor skills ^20^. To investigate whether the observed phenotypic effects of Fingolimod were associated with altered neuronal morphology, we examined neuronal structure and integrity across experimental groups. Cerebellar sections were immunostained with calbindin to label Purkinje cells. Both male and female *Npc1*^mut^ mice exhibited pronounced Purkinje cell neurodegeneration, characterized by neuronal soma degeneration and substantial PC loss (**Fig. 1G-J)**. Notably, Fingolimod treatment preserved Purkinje cell morphology and neuronal soma structure in male *Npc1*^mut^ mice, as evidenced by higher Purkinje cell counts compared with untreated *Npc1*^mut^ animals (**Fig. 1G, H**).

Similar trends toward preserved neuronal soma integrity and reduced Purkinje cell degeneration were also observed in Fingolimod-treated female mice, although these effects were not statistically significant (**Fig. 1I, J**). Gender based effect of Fingolimod has been previously reported in other neurodegenerative diseases ^21^ Overall, our data showed that the observed phenotypical changes and gender-based effect might stem from neuronal protection features. Although the mechanisms underlying this neuroprotective effect is still not clear, the immunomodulatory feature is one of the main proposed mechanisms for its neuroprotective effect ^22,23^.

### Fingolimod Alleviates Neuroinflammation by Targeting both Microglia and Astrocytes

Next, we investigated the effect of Fingolimod on neuroinflammation, a key pathological feature of NPC that contributes significantly to disease progression ^24^. Brain sections were immunostained for IBA1 (ionized calcium-binding adaptor molecule 1) and GFAP (glial fibrillary acidic protein) to assess microglial and astrocytic abundance and activation. Microglial number and morphology were analyzed to evaluate their contribution to disease pathology and to assess potential sex-specific effects of Fingolimod (**Fig. 2A-H**). In line with the previous reports ^14,25^, we observed a significant increase in microglial numbers in both male (**Fig. 2A, B**) and female (**Fig. 2C, D**) *Npc1*^mut^ mice compared to the control groups. Microglia in *Npc1* mutant mice also exhibited marked morphological changes, characterized by a transition from a ramified to a rounded, amoeboid-like phenotype (**Fig. 2E, F**). Quantitative morphometric analysis further revealed a significant reduction in process length and branching complexity, consistent with an activated pro-inflammatory state **(Fig. 2G, H)**. Previous work has demonstrated that loss of NPC1 disrupts cholesterol transport during efferocytosis, leading to expansion of the gastrosome and concomitant adoption of an amoeboid microglial morphology ^25^. In line with these findings, the circular microglia observed in *Npc1*^mut^ mice likely reflect defective cholesterol handling following neuronal engulfment.

**Figure 2.**
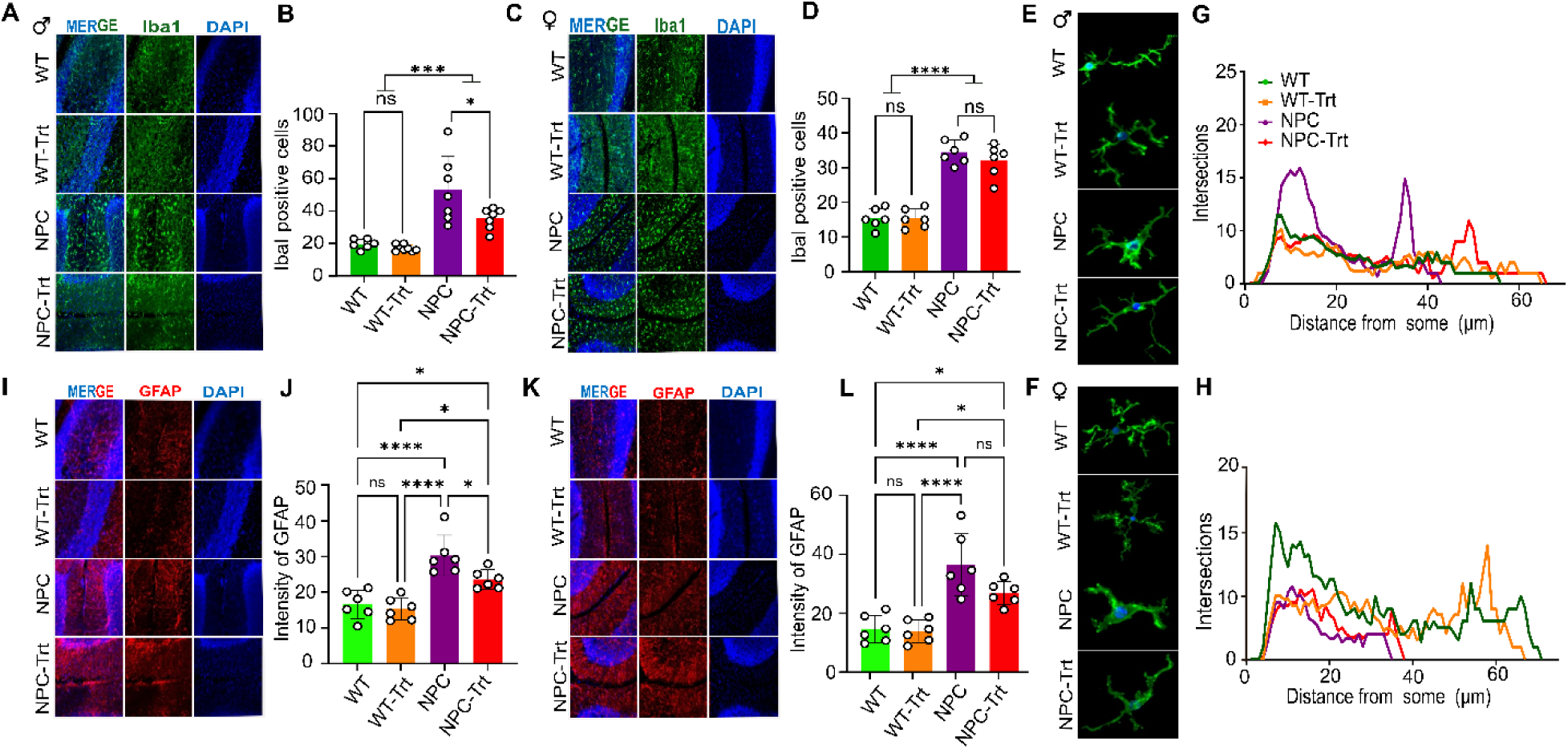
Fingolimod reduces neuroinflammation in a sex-dependent manner in NPC1mut mice. (**A–D**) Confocal images of microglia (Iba1, green) and nuclei (DAPI, blue) in the molecular layer of the cerebellum from male (**A, B**) and female (**C, D**) mice. Experimental groups include wild-type (WT), WT treated with fingolimod (WT-Trt), NPC1mut (NPC), and NPC1mut treated with fingolimod (NPC-Trt). Quantification shows a significant increase in microglial number in NPC1mut mice compared to WT in both sexes. Fingolimod significantly reduced microglial numbers in male NPC1mut mice, but not in females. (**E, F**) Representative images illustrating microglial morphology across groups in male and female mice, respectively. (**G, H**) Sholl analysis of microglial branching complexity. NPC1mut mice show reduced branching compared to WT in both sexes. Fingolimod treatment significantly restored branching complexity in male NPC1mut mice to near WT levels, with no significant effect in females. (**I–L**) Confocal images of astrocytes (GFAP, red) and nuclei (DAPI, blue) in male (**I, J**) and female (**K, L**) mice. Both male and female NPC1mut mice show increased GFAP intensity compared to WT, indicating astrogliosis. Fingolimod significantly reduced astrocyte activation in male NPC1mut mice, but not in females. Confocal images, 20× magnification. Data are mean ± SEM. *p < 0.05, **p < 0.01, ***p < 0.001.

Interestingly, the comparison between untreated and treated *Npc1*^mut^ group with Fingolimod showed a significant effect on decreasing microglia numbers in treated male mice, however no significant effect was observed in female mice (**Fig. 2A-D**). In addition, Fingolimod treatment led to the morphological shifts from an activated, amoeboid form to a surveillant, highly branched (ramified) state in male mice group (**Fig. 2E**). In addition, Sholl analysis of microglial branching in female and male *Npc1*^mut^ mice exhibited a reduction in microglial branching compared to wild-type controls in both sexes. Notably, in male *Npc1*^mut^ mice, Fingolimod treatment significantly restored branching complexity to levels comparable to wild-type mice (**Fig. 2G)**. These findings, together with morphological images, suggest microglial activation in both male and female *Npc1*^mut^ mice, which was significantly ameliorated by Fingolimod treatment in males (**Fig. 2E-H**).

Similarly, an increased GFAP intensity of astrocytes in *Npc1*^mut^ mice compared to control group further indicated a neuroinflammatory state in *Npc1*^mut^ mice **(Fig. 2I-L)**. The observed changes were more drastic in male mice than female mice, indicating a sex-specific severe course of the disease phenotype. Following the treatment, analysis of astrocyte intensity showed the similar sex-specific modulatory effects of Fingolimod. We observed significant reduction in GFAP levels, which corresponds to a decrease in active astrocyte (astrogliosis) levels following treatment in male *Npc1*^mut^ mice (**Fig. 2I, J**) but not in female mice (**Fig. 2K, L**). More importantly, comparison between non-treated and treated control groups showed no significant difference in active microglial and astrocyte levels indicating no toxic effects of the administered dose on healthy young (3-months old) controls of both sexes (**Fig. 2**). In line with our findings, the gender-based effects of Fingolimod also have been reported in MS patients ^[33]^.

### Dysfunctional NPC1 Is Associated with Increased DNA Damage

NPC disease is well known for increased neuroinflammation and loss of motor neurons specifically in cerebellum and particularly Purkinje neurons ^26^, however, we also previously reported increased inflammation and accelerated aging phenotype based on the analyses of frontal cortex from *Npc1*^mut^ mice, indicating NPC1 deficiency systemically affects different parts of the brain.

Although neuroinflammation, lysosomal dysfunction and related pathways (e.g., mTORC1 signaling) have been extensively studied and reported as the main molecular phenotypes ^14,27^, there have not been additional hallmarks reported, associated with the NPC disease. Loss of genomic integrity, driven by accumulated DNA damage and reduced repair capacity, has been proposed as one of the key drivers of aging and neurodegeneration. The aging brain exhibits increased DNA lesions (e.g., double-strand breaks) and impaired repair, leading to cell death or senescence thereby linked to the decline of neuronal functions with age and in neurodegenerative diseases ^28,29^.

Previous reports have demonstrated impaired expression of the DNA sensor cyclic GMP–AMP synthase (cGAS) despite cGAS-independent STING signaling in NPC1-deficient cells ^14^; however, to our knowledge, a systematic assessment of DNA damage in the brains of NPC disease models has not previously been reported. Using γH2AX immunostaining as a marker of DNA double-strand breaks, we detected a substantial increase in DNA damage in the cerebella of both male and female *Npc1*^mut^ mice in comparison to the wild-type controls. Quantification of γH2AX-positive nucleus confirmed a significant elevation in DNA double-strand break burden in *Npc1*^mut^ animals. However, comparison between Fingolimod-treated and untreated *Npc1*^mut^ mice revealed no significant difference in the number of γH2AX-positive nuclei in either sex (**Fig. 3A-D**), indicating that Fingolimod does not modulate DNA damage. To further investigate the DNA damage, we examined γH2AX in primary fibroblasts derived from NPC patient and healthy age-matched control. Analysis of γH2AX immunostaining revealed a similar increase of DSB in NPC patient–derived fibroblasts in comparison to the healthy control cells. Similarly, Fingolimod treatment did not cause any alteration in γH2AX foci number in these cells (**Fig. 3E and F**)

**Figure 3.**
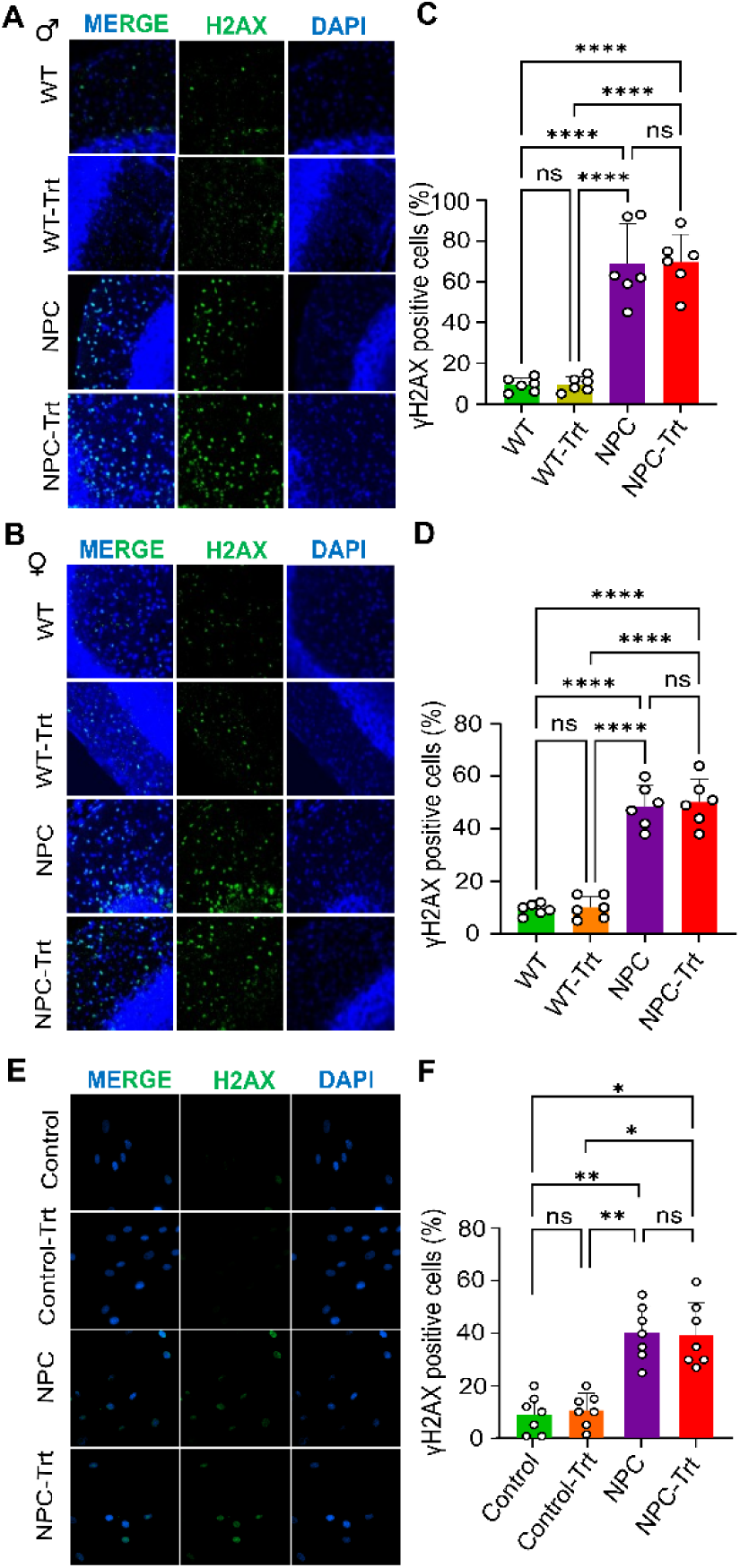
DNA damage is elevated in NPC1mut brain and patient-derived fibroblasts and is not altered by fingolimod treatment. (**A-B**) Representative confocal images of cerebellar sections from male (**A**) and female (**B**) mice showing nuclei (DAPI, blue) and DNA double-strand breaks (γH2AX, green). Experimental groups include wild-type (WT), WT treated with fingolimod (WT-Trt), NPC1mut (NPC), and NPC1mut treated with fingolimod (NPC-Trt). (**C-D**) Quantification of γH2AX-positive nuclei in male (**C**) and female (**D)** mice, respectively. NPC1mut mice exhibit a ∼5–6-fold increase in DNA damage compared to WT controls in both sexes, with a more pronounced effect in males. Fingolimod treatment does not significantly reduce DNA damage in NPC1mut mice in either sex. (**E**) Representative confocal images and quantification of γH2AX staining in human fibroblasts derived from healthy controls (CTRL) and NPC patients. (**F**) NPC fibroblasts show significantly increased DNA damage compared to controls, whereas fingolimod treatment does not alter γH2AX-positive cell numbers. Confocal images, 40× magnification. Data are mean ± SEM. *p < 0.05, **p < 0.01, ***p < 0.001.

It has been shown that DSB-bearing neurons express antiviral and senescence-associated genes and subsequently acquire a senescence-associated secretory phenotype (SASP) that can activate microglia, suggesting that crosstalk between DNA damage and the cGAS–STING pathway contributes to neuroinflammation ^30,31^. In agreement with these findings, we also detected a substantial increase in cGAS expression in the cerebella of both male and female *Npc1*^mut^ mice in comparison to the wild-type controls (**Supplementary Figure S1**). However, in NPC disease, STING activation may also occur independently of cGAS and through non-canonical regulatory mechanisms ^14^. Collectively, these findings suggest that while NPC1 deficiency is associated with increased DNA damage in the cerebellum, Fingolimod does not appear to affect the underlying DNA damage. This dissociation indicates that Fingolimod exerts its anti-inflammatory and potentially neuroprotective effects primarily through modulation of inflammatory signaling pathways rather than by preventing DNA damage itself.

### DNA Double-Strand Breaks and Transcriptional Regulation in NPC Patient-Derived Fibroblasts

Next, we quantified genome-wide DNA DSBs, by using high-resolution DSB mapping/sequencing as previously described ^32^, in both control and NPC fibroblast cells. In addition, to examine whether DSB in NPC cells is associated with gene expression, we also performed RNAseq analyses. DNA break mapping analyses revealed a higher number of DSB clusters sites in NPC fibroblast cells compared to control cells and the number of breaks were enriched in abundantly expressed (higher fpkm) genes in NPC cells (**Fig 4A).** Further analyses of characteristics of these DNA DSBs revealed that they mainly overlapped with regions containing G-quadruplex (G4) structures particularly in NPC cells (**Fig. 4B**). In addition, DNA DSBs were preferentially located at the proximal transcription start sites (TSSs) of both down and up-regulated genes in NPC cells (**Fig. 4C-E**). We further confirmed this by analyzing the DSB enrichment of TSSs on both the down (**Fig. 4F**) and up-regulated (**Fig. 4G**) genes from the RNAseq data, visualized exact break points at single nucleotide resolution for 3 down and 3 up-regulated genes. To further validate the increased G4 signal, we stained NPC cells with G4 structure-specific antibody (BG4). Our analysis revealed increased G4 formation, further confirming the result from DNA break mapping result (**Supplementary Fig. S2**). These findings demonstrated that the structural predisposition of G4 motifs to form energetically stable secondary structures, contributing to genomic instability and gene expression in *NPC1*^mut^ cells, that has never been reported before. Overall, DSB mapping data suggest a possible role of G4 structures in genomic instability and altering gene expression in NPC cells. These results are consistent with published reports that G4s folded in proximal promoter regions are associated either with positive or negative transcriptional regulations ^33,34^.

**Figure 4.**
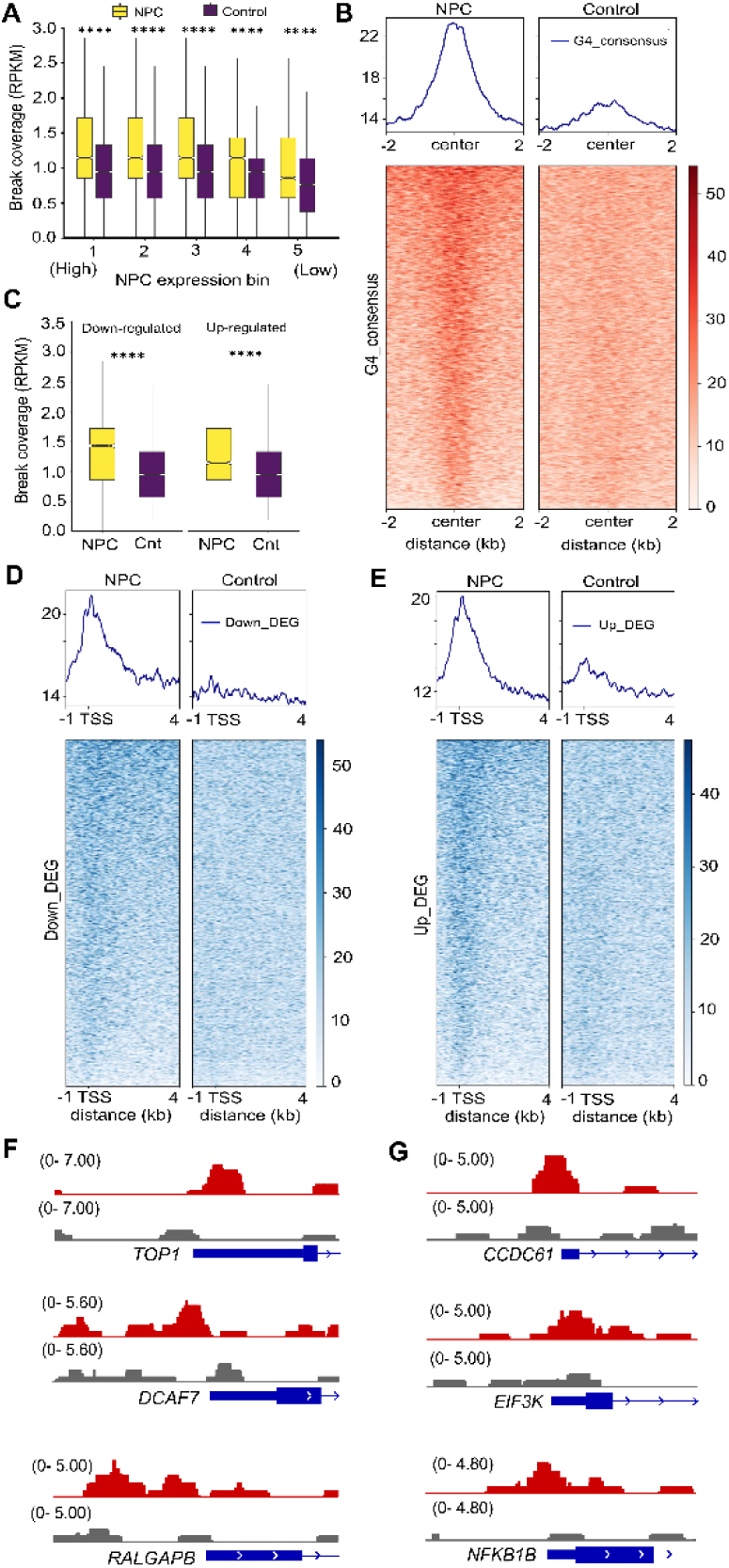
Genome-wide mapping of DNA double-strand breaks reveals enrichment at G-quadruplex (G4) motifs and transcription start sites of differentially expressed genes in NPC fibroblasts. (**A**) Increased DNA double-strand break (DSB) accumulation at transcription start sites (TSS ±250 bp) of differentially expressed genes (DEGs) in NPC fibroblasts compared to healthy controls. Genes were stratified into five equal expression bins based on FPKM values in NPC samples (Bin 1: highest 20% expression; Bin 5: lowest 20%). Boxplots show normalized Break-Seq signal (RPKM) at TSS regions, with NPC samples showing significantly elevated DSB enrichment across all expression bins. Statistical significance was assessed using a paired one-sided Wilcoxon signed-rank test (**** p < 0.0001). **(B**) Genome-wide enrichment of DNA breaks at G-quadruplex (G4) consensus sites and their association with transcriptional regulation. Average profile plots show normalized DSB signal surrounding G4 motifs (±2 kb), demonstrating increased break accumulation in NPC fibroblasts compared with controls. (**C**) Distribution of DNA breaks across G4-associated regions stratified by gene expression changes. DSB enrichment is shown for G4 sites associated with both upregulated and downregulated genes, indicating that G4 structures are broadly linked to transcriptional dysregulation in NPC cells. (**D, E**) Metagene analysis of DNA break distribution at transcription start sites (TSS ±1 kb) of downregulated genes in NPC fibroblasts. Heatmaps and average profiles show increased DSB accumulation at promoter regions in NPC cells compared with controls. Genes are ranked by signal intensity in the NPC condition, and control samples are plotted in the same order to enable direct comparison. Metagene analysis of DNA break distribution at TSS regions of upregulated genes in NPC fibroblasts. Similar to downregulated genes, NPC cells exhibit pronounced enrichment of DSBs at promoter regions, consistent with widespread transcription-associated genomic instability. (**F**) Representative locus-specific Break-Seq profiles at single-nucleotide resolution for selected downregulated (TOP1, DCAF7, RALGAPB, PTGIS) and (**G**) upregulated genes (CCDC61, EIF3K, NFKB1B, SMG9). Tracks show normalized DSB read density in NPC (red) and control (gray) fibroblasts. Despite higher total sequencing depth in controls, NPC samples display markedly increased DSB accumulation at transcription start sites. Gene models are shown below each track, with exon structure and transcriptional direction indicated.

Finally, to define the altered transcriptomic consequences of NPC1 deficiency in a disease context, we comprehensively analyzed our RNAseq data (**Supplementary File S1**). Comparative analysis identified 9,669 differentially expressed genes (DEGs) at a stringent false discovery rate threshold (Benjamini–Hochberg adjusted p-value < 0.05), comprising 4,867 upregulated and 4,802 downregulated transcripts in NPC relative to control cells. Applying an additional effect-size criterion (|log_2_FC| > 1.5), resulted in 3,292 robustly up and 3,723 downregulated genes (**Fig. 5A, Supplementary Fig. S3)**. The magnitude of significant changes of transcriptional response indicated that NPC1 dysfunction imposes a profound, multi-pathway cellular reprogramming that extends far beyond a focal lipid-storage perturbation, including immune response, mitochondrial function, mTORC signaling and unfolded protein response (**Fig. 5B)**.

**Figure 5.**
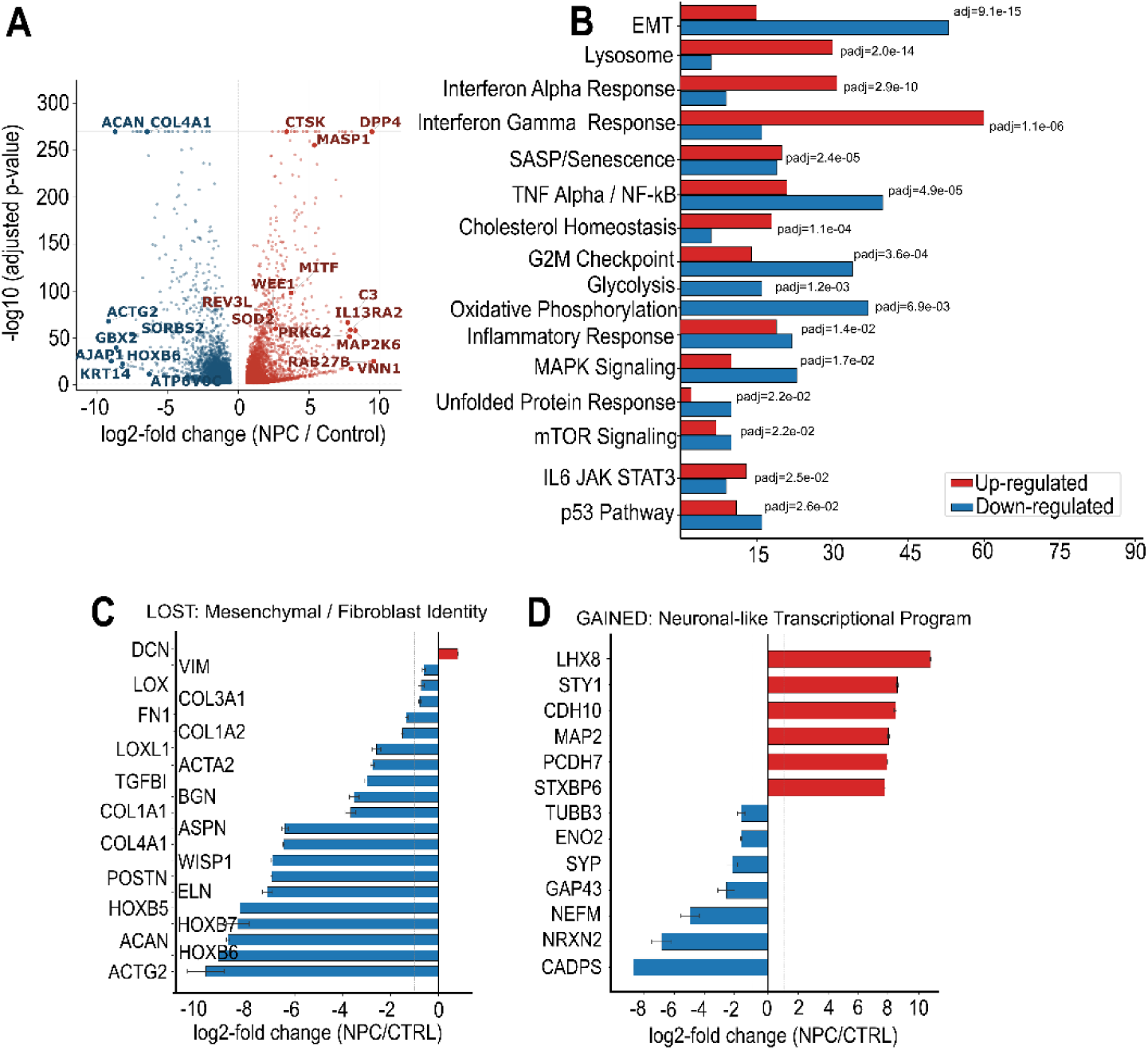
NPC1 deficiency induces global transcriptomic reprogramming characterized by lysosomal activation, cholesterol dysregulation, inflammatory signaling, and loss of fibroblast identity. **(A)** Differential gene expression analysis of NPC patient-derived fibroblasts compared with healthy controls. Volcano plot summar identifies 9,669 differentially expressed genes (DEGs) at FDR < 0.05, including 4,867 upregulated and 4,80 downregulated transcripts. Applying an additional effect-size cutoff (|log2FC| > 1.5) identifies 3,292 strongly upregulated and 3,723 strongly downregulated genes, reflecting extensive transcriptomic remodeling in NPC cells. **(B)** KEGG pathway over-representation analysis of upregulated genes showing significant enrichment of lysosomal and lipid metabolism pathways. Key induced genes include lysosomal hydrolases (e.g., cathepsins), NPC2, LAMP2, and SGMS1, indicating enhanced lysosomal biogenesis and compensatory cholesterol trafficking responses. **(C)** Suppression of fibroblast and mesenchymal identity programs in NPC fibroblasts. Heatmap and enrichment analysis of the Hallmark epithelial–mesenchymal transition (EMT) gene set show significant downregulation of structural ECM genes (COL1A1, COL4A1, ELN, POSTN), cytoskeletal regulators (ACTA2, ACTG2), and HOX-cluster transcription factors (HOXB5, HOXB6, HOXB7), indicating loss of fibroblast identity. **(D)** Aberrant induction of neuronal-lineage gene expression in NPC fibroblasts. Bar plots/heatmap illustrate strong upregulation of neuronal markers including MAP2, SYT1, LHX8, CDH10, and PCDH7, along with significant enrichment of neuronal differentiation gene sets, despite absence of terminal neuronal maturation markers (e.g., NEFL, TUBB3), suggesting partial and non-functional neuronal transcriptional reprogramming.

As expected, pathway over-representation analysis revealed highly significant enrichment of KEGG lysosomal pathway genes among up- and downregulated DEGs (**Fig. 5B**). Lysosomal cysteine cathepsins were among the most consistently induced transcripts (**Supplementary Fig. S4 and Supplementary File S1**). *NPC2*, the soluble intralysosomal cholesterol-shuttle protein whose function is complementary to *NPC1*, was significantly upregulated (log_2_FC = +1.55, adj. p = 2.6 × 10^−41^), consistent with a compensatory attempt to sustain cholesterol egress in the absence of functional NPC1. The lysosomal membrane glycoprotein coding *LAMP2* and sphingomyelin synthase *SGMS1* genes were likewise elevated, together pointing to coordinated expansion of lysosomal biogenesis capacity ^[35–38]^.

Another significant finding was a marked dissociation between the two principal transcriptional regulators of the CLEAR network ^35–37^. *TFEB*, the canonical master regulator of lysosomal biogenesis, was significantly downregulated (log_2_FC = −1.22, adj. p = 1.1 × 10^−6^), as was its paralogue *TFE3* (−0.61, adj. p = 1.8 × 10^−4^). In stark contrast, *MITF* a member of the same MiT/TFE family that binds identical CLEAR-box promoter elements was highly induced (log_2_FC = +3.71, adj. p = 1.5 × 10^−98^) (**Supplementary Fig. S4**). These data indicate that the lysosomal gene program in NPC fibroblasts may be driven primarily through MITF rather than TFEB, a distinction with direct relevance for therapeutic strategies premised on pharmacological TFEB activation ^37^.

Despite the well-established accumulation of unesterified cholesterol within the NPC lysosomal compartment, multiple cholesterol biosynthesis genes were paradoxically upregulated in patient fibroblasts, including *HMGCR* (+0.78, adj. p = 1.7 × 10^−14^), *HMGCS1* (+0.72), SQLE (+0.85), *FDFT1* (+0.73), and *DHCR24* (+2.83, adj. p = 6.6 × 10^−107^). Simultaneously, the ATP-binding cassette transporter *ABCA1* the primary mediator of cholesterol efflux to apolipoprotein A was among the most significantly induced genes in the dataset (log_2_FC = +2.08, adj. p = 2.2 × 10^−91^), and the lipid transfer protein *OSBPL1A* (+1.36, adj. p = 5.8 × 10^−48^) was similarly elevated, suggesting a compensatory attempt to redistribute cholesterol away from the sequestered lysosomal pool (**Supplementary Fig. S4 and Supplementary File S1)**.

Systematic analyses of inflammatory signaling genes revealed a multi-layered, non-canonical inflammatory state that was dominated by a broadly activated interferon-stimulated gene (ISG) program and JAK-STAT signaling (**Supplementary Fig. S5**). Twenty-seven of 36 curated ISGs were significantly upregulated (hypergeometric p = 1.3 × 10^−14^), including the 2′-5′ oligoadenylate synthases *OAS2*, *OAS1*, the antiviral GTPases GBP2 and *GBP4*, *HERC5*, *IFI44L*, and the *IFIT* family members *IFIT1*, *IFIT2*, and *IFIT3* (**Supplementary Fig. S5F**). In the absence of an exogenous viral trigger, this ISG induction may reflect activation of the cGAS–STING innate DNA-sensing pathway by cytosolic nucleic acids. Supporting this interpretation, the cytosolic nucleic acid sensors *IFIH1/MDA5* (log_2_FC = +3.49, adj. p = 6.3 × 10^−34^) and *DDX58/RIG-I* (log_2_FC = +2.44, adj. p = 8.3 × 10^−45^) were significantly induced (**Supplementary Fig. S5 and Supplementary File S1)**.

Finally, transcriptome-wide enrichment analysis revealed profound suppression of mesenchymal and fibroblast identity programs in NPC fibroblasts. Genes of the Hallmark Epithelial–Mesenchymal Transition program were significantly downregulated (28 of 40 overlapping genes; hypergeometric adj.p = 9.18 × 10^−15^), encompassing contractile cytoskeletal proteins *ACTG2* (log_2_FC = −9.23, adj. p = 2.8 × 10^−68^) and *ACTA2* (−2.76, adj. p = 2.7 × 10^−157^), extracellular matrix structural proteins elastin *ELN* (log_2_FC = −7.17), periostin *POSTN* (log_2_FC = −7.00), *COL4A1* (log_2_FC = −6.49), and *COL1A1* (−3.68), and genes coding for matrix cross-linking enzymes *LOX* and *LOXL1* (**Fig. 5C)**. The HOX-cluster transcription factors *HOXB5*, *HOXB6*, and *HOXB7*, which collectively define fibroblast positional identity, were among the most severely suppressed transcripts in the dataset. Paradoxically, this programmatic loss of fibroblast identity was accompanied by significant induction of neuronal-lineage markers (**Fig. 5D)**. The dendritic arbour protein coding gene *MAP2* was induced by over 185-fold (log_2_FC = +7.53, adj. p = 1.1 × 10^−269^), synaptotagmin *SYT1* by over 220-fold (+7.81, adj. p = 4.8 × 10^−56^), and the neural transcription factor *LHX8* normally restricted to the developing basal ganglia by over 600-fold (log_2_FC = +9.24, adj. p = 9.3 × 10^−25^). Neuronal-cadherin *CDH10* (log_2_FC = +7.48) and protocadherin *PCDH7* (log_2_FC = +7.24) were similarly elevated, and neuronal differentiation marker genes were significantly over-represented among upregulated DEGs (hypergeometric p = 2.5 × 10^−3^). Critically, terminal neuronal maturation markers including *NEFL*, *TUBB3*, and classical synaptic-vesicle proteins were not altered significantly, arguing against genuine neuronal trans-differentiation and instead implicating aberrant, partial activation of a neural transcriptional program superimposed on a de-differentiated cellular background. This phenomenon has been reported in Hodgkin lymphoma, where neighbouring B cells lose core B-cell identity genes due to microenvironmental influence from Hodgkin and Reed–Sternberg cells ^[47]^.

Taken together, these transcriptomic findings delineate NPC1 deficiency as driving a multidimensional cellular crisis encompassing lysosomal biogenesis rewiring, cholesterol-sensing paradox, autophagy blockade, adaptive ER stress, near-complete fibroblast dedifferentiation with aberrant neural programming, and a multi-pathway inflammatory state that collectively redefines the molecular landscape of NPC disease in non-neuronal cells.

### Fingolimod Induces a Disease-Specific Transcriptional Rescue Response in NPC Patient Fibroblasts

To evaluate the transcriptional consequences of Fingolimod treatment in NPC1-deficient cells, we performed genome-wide RNA sequencing of Fingolimod-treated and untreated NPC patient fibroblasts and healthy control fibroblasts. The effect was analyzed based on the genes, whose NPC disease expression signature established in the untreated NPC vs control comparison was significantly reversed in direction by Fingolimod treatment assessed through the pairwise comparison of treated versus untreated NPC fibroblasts (NPC-T vs NPC). This approach separates the drug’s disease-specific therapeutic action from its baseline cellular effects in healthy cells. Fingolimod induced 1,959 significant transcriptional changes in control fibroblasts (1,114 upregulated, 845 downregulated; Benjamini–Hochberg adjusted p < 0.05, **Figure 6A and Supplementary File S1**) and 5,137 changes in NPC fibroblasts (2,706 upregulated, 2,431 downregulated; **Figure 6B and Supplementary File S1**), with the substantially larger NPC response indicating amplified pharmacological sensitivity in the disease context. To quantify the disease-correcting capacity of Fingolimod, we performed a gene-level rescue analysis and found that 2,805 of the 7,016 NPC dysregulated genes (40.0%) were corrected back toward control-level expression by Fingolimod treatment comprising 1,530 genes that were downregulated in NPC and restored upward, and 1,275 genes that were upregulated in NPC and corrected downward (**Supplementary Fig. S6A-D**).

**Figure 6.**
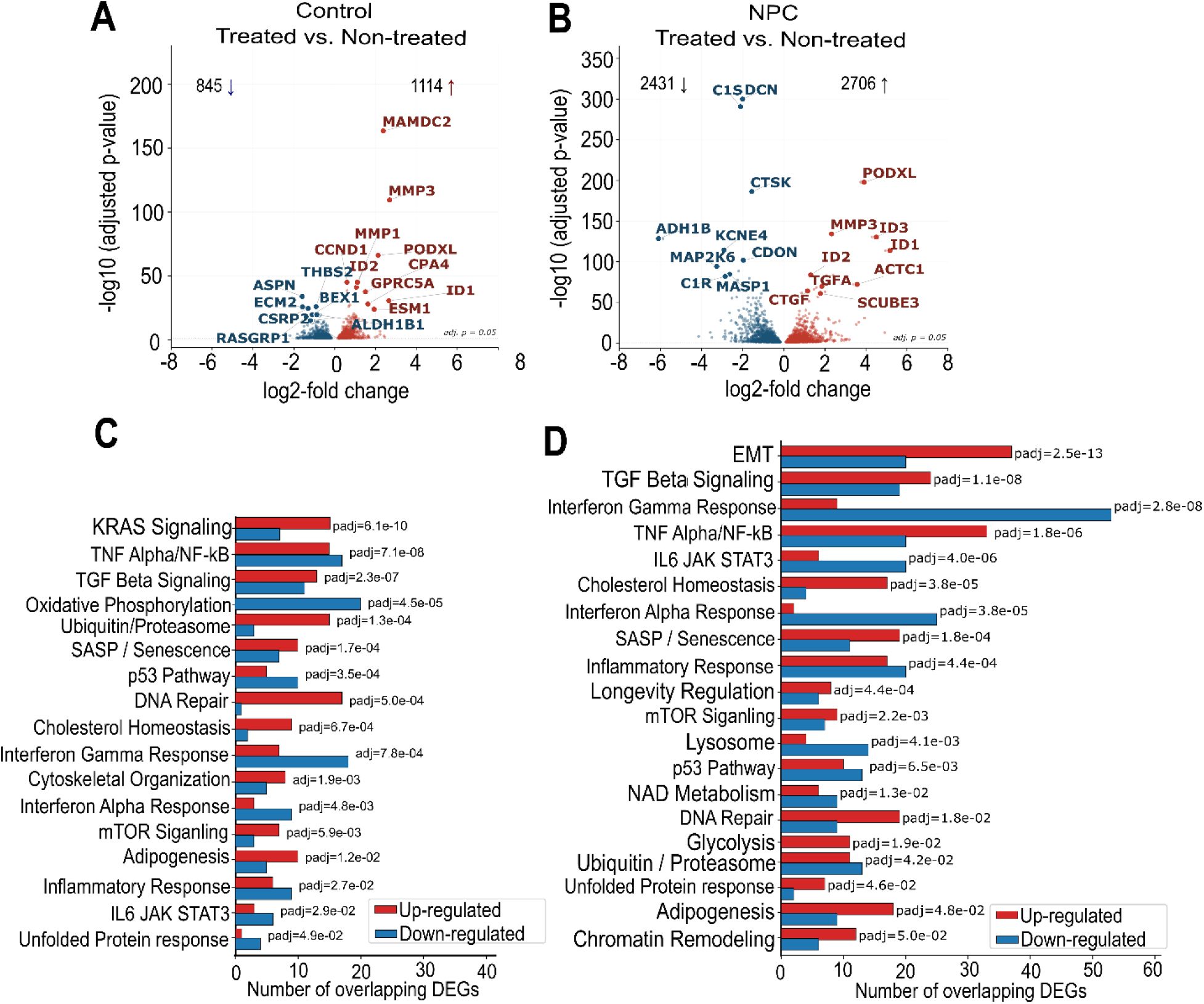
Fingolimod induces a disease-specific transcriptional rescue program in NPC patient-derived fibroblasts. Genome-wide RNA sequencing was performed on untreated and Fingolimod-treated NPC patient fibroblasts and healthy control fibroblasts. (**A, B**) Differential expression analysis showing the number of Fingolimod-responsive genes in control fibroblasts (**A**; 1,959 genes: 1,114 upregulated, 845 downregulated) an NPC fibroblasts (**B**; 5,137 genes: 2,706 upregulated, 2,431 downregulated), indicating a markedly enhanced transcriptional response in the disease context. (**C, D**) Pathway-level enrichment analysis demonstrating preferential rescue of lysosomal, cholesterol homeostasis, complement, inflammatory, and oxidative stress-related gene programs in NPC cells following Fingolimod treatment.

Although several key pathways commonly targeted in both control and NPC groups, such as cholesterol homeostasis, DNA repair, inflammatory responses and unfolded protein responses, Fingolimod exerted its most functionally significant rescue in the lysosomal, cholesterol homeostasis, and complement programs that define the core NPC disease state **(Fig. 6C, D)**.

Inflammatory signaling was significantly attenuated by Fingolimod in a mechanistically coherent manner. The cytokines *IL-6* (adj. p = 2.3 × 10^−27^) and CCL2 (adj. p = 1.7 × 10^−39^) were significantly reduced. The interferon-stimulated gene (ISG) program was partially dampened, with *IFIT1*, *OAS2*, and *DPP4* showing significant reductions in treated NPC cells, though the ISG signature remained substantially elevated above control levels. Extracellular matrix gene expression was also partially rescued: elastin *ELN*, *COL4A1*, and periostin *POSTN* were significantly induced in treated NPC cells **(Supplementary Fig. S6B)**, consistent with partial restoration of fibroblast mesenchymal identity. Overall, pathway-level over-representation analysis confirmed significant enrichment of lysosomal, EMT-restoration, complement, and anti-inflammatory genes among Fingolimod-responsive NPC transcripts (**Supplementary File S1**).

Finally, considering the increased DNA damage signal from the mice cerebellum and patient-derived NPC cells, we examined the expression pattern of DNA repair and genome stability genes. Transcriptome-wide screening of 126 curated DNA repair and genome stability genes across all four conditions revealed a coherent and previously unreported dimension of NPC pathology: a shift from high-fidelity repair toward error-prone DNA damage tolerance that is substantially reversed by Fingolimod **(Supplementary Fig. S7 and Supplementary File S1)**. In untreated NPC fibroblasts, the G_2_/M checkpoint kinase *WEE1* was among the most strongly induced transcripts in the dataset (disease log_2_FC = +2.43), alongside the DNA damage checkpoint kinase *CHEK2*, the *ATM* kinase, and the p53 regulator *MDM2*, indicating that untreated NPC cells are under chronic replication stress and arrest before mitosis. Concurrently, multiple homologous recombination (HR) pathway components responsible for high-fidelity double-strand break resolution were significantly suppressed: *RAD54L*, *RECQL4*, *EME2*, *SLX4*, and *MUS81*, along with the Fanconi anemia genes *FANCA*, *FANCE*, and *FANCG*. The genes coding for base excision repair (BER) scaffold protein *XRCC1*, the mitochondrial polymerase *POLG*, *MUTYH*, and *FEN1* were all downregulated, consistent with impaired repair of the oxidative DNA lesions ^38^. Most strikingly, the translesion synthesis (TLS) polymerases *REV3L* the catalytic subunit of the highly mutagenic Pol ζ complex and *POLI*, *POLK*, and *ZRANB3* were induced, indicating that NPC cells bypass unrepaired DNA lesions via error-prone mechanisms when the HR and BER pathways are compromised. Together, these findings define a chronic DNA damage tolerance phenotype in NPC fibroblasts: BER and HR suppressed, TLS polymerases elevated, and checkpoint constitutively activated a pattern mechanistically linked to the lysosomal ROS burden identified in untreated NPC (**Supplementary Fig. S8)**.

Fingolimod treatment in NPC cells substantially corrected this genome instability program in an NPC-specific manner (53 of 64 treatment effects absent in treated healthy control cells). *WEE1* was significantly reduced (treatment log_2_FC = −1.12, adj. p = 1.0 × 10^−17^) and *CHEK2* (adj. p = 2.8 × 10^−7^) was downregulated, consistent with release of the G_2_/M arrest as DNA damage burden decreases. The mutagenic TLS polymerases were significantly suppressed: REV3L (adj. p = 2.7 × 10^−23^), *POLI*, *POLK*, and *ZRANB3* were all downregulated in treated NPC cells, whilst HR pathway components *RAD54L*, *EME2*, *SLX4*, *RECQL4*, and *FANCA* were restored indicating a return toward high-fidelity DSB repair. The BER scaffold *XRCC1*, *POLG*, *LIG1*, and *MUTYH* were partially reinstated. Notably, Fingolimod exerted a distinct and non-overlapping effect in healthy control cells, where it induced *BRCA1*, *BRCA2*, *BLM*, and *PARP1* without the TLS suppression observed in NPC cells, confirming that the genomic-repair rescue in NPC is a true therapeutic response to disease-specific DNA damage rather than a general pharmacological effect (**Supplementary Fig. S8B)**.

Taken together, these data demonstrate that Fingolimod produces a stratified, functionally coherent partial rescue of the NPC transcriptome, preferentially normalizing the metabolic and inflammatory consequences of lysosomal dysfunction, complement activation, DNA damage response while leaving the fundamental cell-state reprogramming and primary innate immune sensor induction largely intact. This pattern is consistent with Fingolimod’s mechanism as a sphingosine-1-phosphate receptor modulator with secondary effects on ceramide and sphingolipid metabolism ^39^,and indicates that complementary strategies targeting the upstream cell-identity crisis and TLR-driven innate sensing would be required for complete transcriptional normalization in NPC.

### Fingolimod Facilitates Lysosomal Cholesterol Trafficking and Modulates STING Signaling Through LAMP1 Regulation

Next, based on previous evidence implicating STING signaling in NPC-associated neuroinflammation ^14^, and our immunofluorescence findings from Npc1^mut^ mice brain sections demonstrating reduced inflammation following Fingolimod treatment, we investigated whether Fingolimod alters phosphorylated STING levels as a potential mechanism underlying its anti-inflammatory effects. Western blot analysis revealed that Fingolimod increased phospho-STING levels in control cells, whereas it reduced phospho-STING levels in NPC patient-derived fibroblasts **(Fig. 7A)**. These findings suggest that Fingolimod modulates STING signaling in a context-dependent manner influenced by the basal inflammatory and lysosomal state of the cell. In control cells, Fingolimod may enhance lysosomal function through STING activation, which has been reported to support proper lysosomal activity ^40^. In contrast, in NPC cells, where STING signaling is chronically elevated and associated with persistent inflammation ^14^, Fingolimod suppresses STING activity, potentially alleviating inflammatory stress. Although phosphorylated STING protein levels were markedly increased in NPC cells and significantly reduced following Fingolimod treatment, analysis of RNA sequencing data revealed an inverse pattern at the transcript level. We found that STING (*TMEM173*) mRNA expression was significantly (adj. p= 1.68 × 10^−7^) decreased in NPC cells compared with healthy controls, and this reduction was not altered by Fingolimod treatment **(Fig. 7B).** Thus, Fingolimod altered active STING protein abundance without affecting STING transcription, suggesting that STING accumulation in NPC cells is regulated primarily through post-transcriptional mechanisms. The reduced STING mRNA levels observed in NPC cells may represent a compensatory transcriptional response aimed at limiting excessive STING signaling associated with chronic protein accumulation. However, this response appears insufficient to normalize STING protein levels, indicating dysregulated protein turnover or impaired degradation pathways, potentially involving lysosomal or proteasomal dysfunction. Consistent with this interpretation, Fingolimod reduced STING protein abundance without altering STING mRNA expression. Given emerging evidence linking STING signaling to aging and longevity, including reports demonstrating that STING deficiency significantly shortens murine lifespan ^41^, we next investigated whether dysregulated STING signaling in NPC cells is associated with senescence-related alterations. To address this, additional western blot analyses were performed to assess p21 levels, a marker of cellular senescence, and to evaluate the effect of Fingolimod on these aging-associated changes. Consistent with our previous findings demonstrating accelerated brain aging in the NPC mouse model ^5^, NPC cells exhibited elevated p21 levels. Notably, Fingolimod treatment reduced p21 expression (**Fig. 7A**), suggesting that modulation of aberrant STING signaling may contribute to the attenuation of senescence-associated phenotypes in NPC cells.

**Figure 7.**
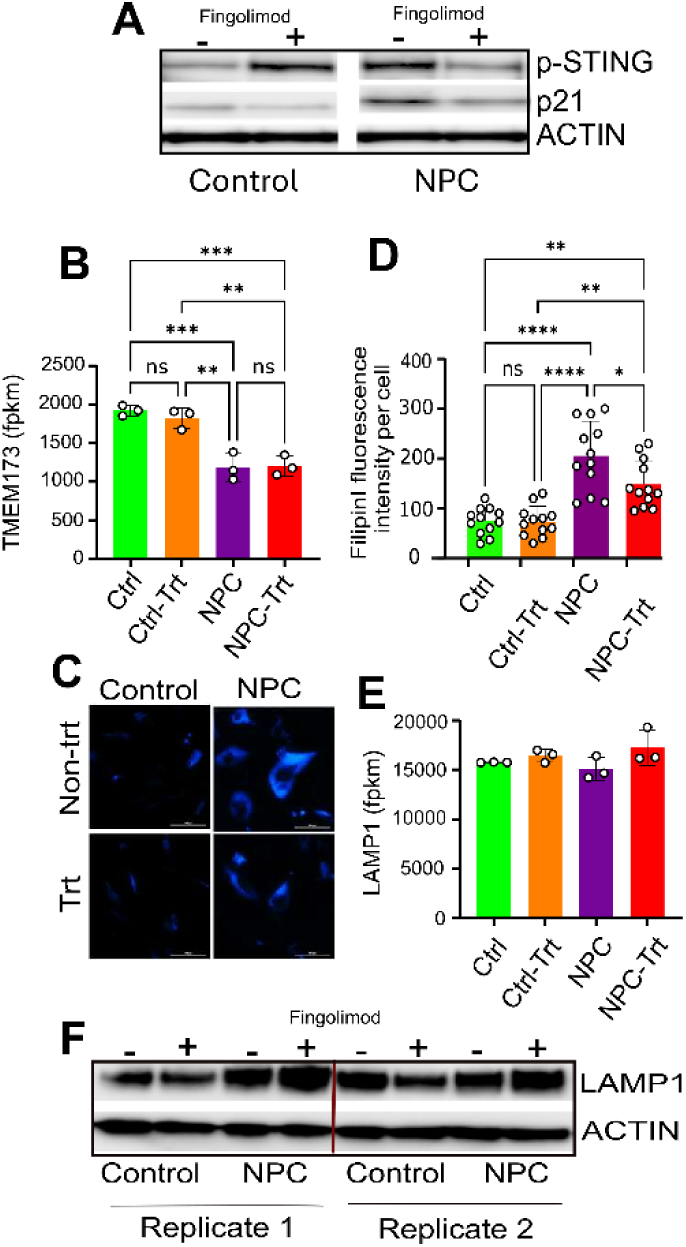
Fingolimod modulates lysosomal cholesterol trafficking and STING signaling via LAMP1-dependent lysosomal adaptation in NPC cells. (**A**) Western blot analysis of phosphorylated STING (p-STING) and p21 in healthy control (CTRL) and NPC patient-derived fibroblasts with or without fingolimod treatment. NPC fibroblasts exhibit elevated p-STING and increased p21 levels, indicating chronic STING activation and senescence-associated signaling. Fingolimod reduces p-STING and p21 levels in NPC cells, while increasing p-STING in control cells, indicating context-dependent regulation. (**B**) RNA-seq analysis of STING (TMEM173) mRNA expression in CTRL and NPC fibroblasts. STING transcript levels are significantly reduced in NPC cells compare with controls and remain unchanged upon fingolimod treatment, indicating post-transcriptional regulation of STING protein abundance. (**C, D**) Representative images and quantification of Filipin III staining in fibroblasts from CTRL and NPC patients. NPC fibroblasts show marked accumulation of cholesterol compared with controls. Fingolimod treatment significantly reduces intracellular cholesterol accumulation in NPC cells. (**E**) RNA-seq analysis of LAMP1 mRNA expression in CTRL and NPC fibroblasts. LAMP1 transcript levels are not significantly altered in NPC cells compared with controls and remain unchanged upon fingolimod treatment, indicating post-transcriptional regulation of LAMP1 protein abundance. (**F**) Two biological replicates of Western blot analysis of lysosome-associated membrane protein 1 (LAMP1) in CTRL and NPC fibroblasts. NPC cells show increased LAMP1 expression compared with controls, and fingolimod further elevates LAMP1 protein levels, suggesting enhanced lysosomal capacity and adaptation. Data are representative of at least three independent experiments and presented as mean ± SEM where applicable. Statistical significance: *p < 0.05, **p < 0.01, ***p < 0.001.

Then, to determine whether Fingolimod modulates lysosomal cholesterol accumulation, a process implicated in aberrant STING activity, autophagic flux and oxidative stress ^[50–52]^, we assessed cholesterol levels in patient-derived NPC fibroblasts using Filipin III staining **(Fig. 7C)**. Consistent with the established pathology of NPC disease, NPC fibroblasts exhibited markedly elevated Filipin III fluorescence compared with healthy controls, indicating substantial accumulation of unesterified cholesterol. Fingolimod treatment significantly reduced Filipin III staining intensity in NPC fibroblasts, suggesting improved intracellular cholesterol handling and lysosomal recovery (**Fig. 7D**). Next, to explore potential mechanisms underlying Fingolimod-mediated cholesterol reduction, we examined lysosome-associated membrane protein 1 (LAMP1), a key structural and functional component of the lysosomal membrane involved in lysosomal integrity and lipid trafficking. Our transcriptome data showed no expression difference between control and NPC cells and Fingolimod treatment also did not alter the mRNA abundance across these samples (**Fig. 7E**). However, in line with previous reports describing variable LAMP1 expression across NPC models depending on cell type and experimental context ^[21^,^53]^, we observed elevated LAMP1 protein levels in NPC patient-derived fibroblasts relative to controls, potentially reflecting a compensatory response at protein level to impaired cholesterol export caused by NPC1 deficiency. Notably, Fingolimod treatment further increased LAMP1 protein abundance (**Fig. 7F**), suggesting that enhanced lysosomal membrane capacity or lysosomal adaptation may contribute to the observed reduction in cholesterol accumulation. Interestingly, a similar mechanism has been proposed for 2-hydroxypropyl-β-cyclodextrin (HPβCD) in rescuing cholesterol accumulation in NPC1-deficient cells ^[53^,^54]^. Altogether, our findings support a model in which Fingolimod promotes lysosomal adaptation and cholesterol clearance, thereby attenuating pathological STING activation and senescence-associated alterations in NPC cells. These results identify lysosomal-STING crosstalk as a potential mechanism underlying the therapeutic effects of Fingolimod in NPC disease.

### Fingolimod Attenuates Mitochondrial Dysfunction

It was reported by several studies that primary lysosomal defect in NPC1 mutant fibroblasts is accompanied by deregulation of the organization and function of the mitochondrial network, which contributes to disease progression ^42,43^. Additionally, previous findings from cultured neuronal cells and frontal cortex and cerebellum from treated mice suggest that Fingolimod may exert its protective effects in part by inducing metabolic reprogramming and directly enhancing mitochondrial function, particularly under oxidative stress ^8,44^. This prompted us to investigate the effect of Fingolimod treatment on mitochondrial properties.

To investigate the impact on mitochondrial function, we assessed mitochondrial membrane potential (ΔΨm), morphology, and mitochondrial DNA (mtDNA) localization in NPC patient-derived fibroblasts compared to healthy controls, with and without Fingolimod treatment. Super-resolution live-cell confocal imaging was performed to assess mitochondrial phenotype (**Fig. 8A**). Notably, mitochondrial morphology appeared broadly similar across all four conditions: control and NPC fibroblasts displayed elongated, reticular mitochondrial networks, indicating that the gross structural organization of the mitochondrial network is largely preserved in NPC cells. Despite this morphological similarity, however, two striking functional differences were evident. First, CMXRos intensity was increased in untreated NPC fibroblasts compared to controls, reflecting elevated ΔΨm. Quantification of ΔΨm confirmed these observations that untreated NPC fibroblasts exhibited a significant elevation in ΔΨm (p < 0.0001), indicating a mitochondrial hyperpolarization **(Fig. 8B**). Treatment with FTY720 significantly reduced ΔΨm in NPC fibroblasts (p < 0.001), bringing membrane potential back down to levels of untreated controls. Fingolimod treatment also did not significantly alter ΔΨm in control fibroblasts, indicating that the normalizing effect is specific to the NPC disease state rather than a generalized suppression of mitochondrial function **(Fig. 8B**).

**Figure 8.**
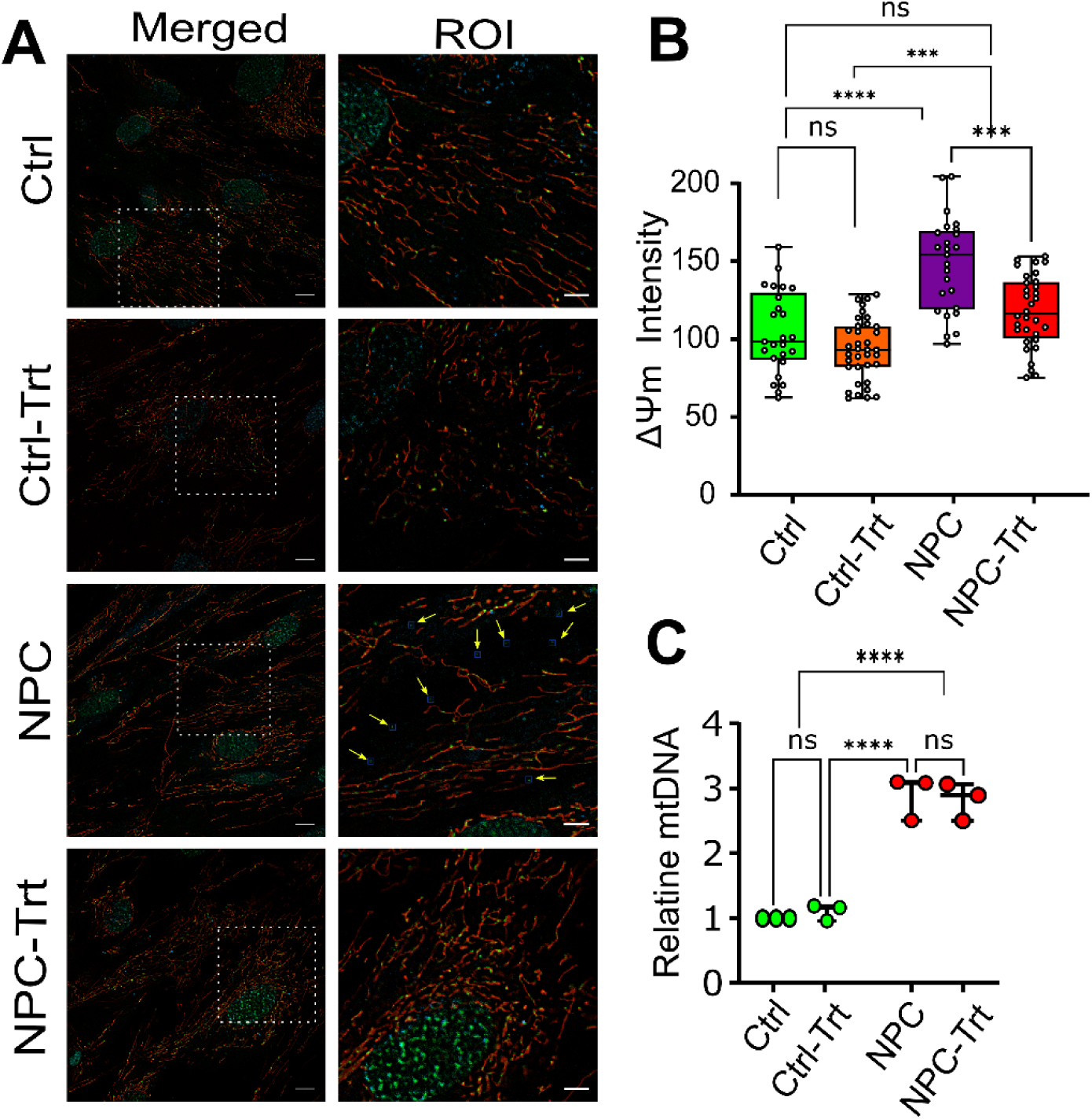
Fingolimod attenuates mitochondrial dysfunction in NPC cells. **(A)** Super-resolution confocal imaging of mitochondrial structure and function in control, NPC, and Fingolimod-treated fibroblasts. All groups exhibited broadly preserved elongated mitochondrial networks, indicating no gross structural disruption. Visualization of mitochondrial DNA (mtDNA) distribution using PicoGreen staining revealed extramitochondrial mtDNA leakage in untreated NPC fibroblasts, which was absent in both control and Fingolimod-treated NPC cells, shown with the yellow arrow. (**B**) NPC fibroblasts displayed increased mitochondrial membrane potential (ΔΨm), as measured by MitoTracker CMXRos fluorescence, which was normalized following Fingolimod treatment without affecting control cells. Quantification of ΔΨm showing significant mitochondrial hyperpolarization in untreated NPC fibroblasts compared to controls, and restoration toward physiological levels following Fingolimod treatment. (**C**) Analysis of relative mitochondrial DNA (mtDNA) copy number revealed significant ∼3-fold increase in mtDNA abundance in NPC fibroblasts relative to controls. These data represent at least three independent experiments and presented as mean ± SEM where applicable. Statistical significance: *p < 0.05, **p < 0.01, ***p < 0.001, ****p < 0.0001.

Second, PicoGreen-labeled mtDNA puncta were clearly visible outside the MitoTracker-defined mitochondrial boundaries in untreated NPC cells (**Fig. 8A**, yellow arrows). These extramitochondrial mtDNA foci were not observed in control fibroblasts (either untreated or Fingolimod-treated) or in Fingolimod-treated NPC cells, suggesting that mtDNA displacement from the mitochondrial network is a specific consequence of the NPC disease state that is reversible upon Fingolimod treatment. Given the higher number of mtDNA foci observed by imaging, we next used transcriptome data to determine whether this reflected a change in overall mitochondrial genome abundance. NPC fibroblasts exhibited an approximately 3-fold increase (p < 0.0001) in relative mtDNA copy number compared with control cells (**Fig. 8C**). Notably, this increase in mtDNA copy number was not significantly affected by Fingolimod treatment in either NPC or control cells (**Fig. 8C**), indicating that Fingolimod corrects mtDNA mislocalization without altering total mitochondrial genome copy number, a dissociation suggesting these two phenotypes arise through distinct mechanisms. The elongated mitochondrial morphology and mitochondrial membrane hyperpolarization observed in NPC fibroblasts are consistent with recent findings in Ncr1-deficient yeast, the yeast homolog of NPC1^45^. Notably, however, Ncr1-deficient yeast cells also exhibited reduced mitochondrial DNA content, indicating a potential divergence in mitochondrial genome maintenance between yeast and mammalian NPC models.

Overall, our data suggests that Fingolimod may exert its protective effects in part by restoring mitochondrial function.

### Fingolimod Promotes Cellular and Organismal Longevity

Given the observed modulatory effects of Fingolimod on common hallmarks of aging ^46^; inflammation, senescence and mitochondrial dysfunction, we investigated whether Fingolimod directly modulates aging. First, to determine whether NPC1 deficiency drives accelerated cellular aging and whether Fingolimod treatment attenuates this phenotype, we applied a transcriptome-based aging clock to the RNA-seq expression profiles **(Fig. 9A)**. The aging score was computed per sample using 100 curated aging-associated genes derived from four published resources, such that a higher score reflects a transcriptome more closely resembling an aged cellular state (see methods). Healthy control fibroblasts produced a mean aging score of −0.46 (± 0.01 SEM, n = 3), reflecting the expected young basal state. Untreated NPC patient fibroblasts exhibited a dramatically elevated mean aging score of +0.61 ± 0.03, representing an increase of +1.08 units relative to controls (Welch t-test: t = 34.24, p = 1.1 × 10^−4^), consistent with transcriptome-wide acceleration of cellular aging driven by NPC1 deficiency (**Fig. 9A**). Fingolimod treatment substantially reversed the NPC aging transcriptome. The mean aging score in treated NPC cells was +0.06 ± 0.03, representing a reduction of −0.55 units relative to untreated NPC (Welch t-test: t = 12.21, p = 3.1 × 10^−4^), corresponding to reversal of the NPC-associated aging acceleration. Nonetheless, Fingolimod treated NPC score (+0.06) remained significantly above the healthy control score (−0.46; Welch t-test: t = 14.16, p = 1.9 × 10^−3^), indicating that Fingolimod only partially decelerates cellular aging that the residual accelerated aging persists after treatment. Notably, Fingolimod treatment of healthy control cells also reduced the aging score (CTRL + Fingolimod = −0.84 ± 0.04; change of −0.38 units; p = 8.9 × 10^−3^), suggesting a modest transcriptional rejuvenating effect of Fingolimod in non-diseased fibroblasts, consistent with its known anti-inflammatory effect **(Fig. 9A)**. Taken together, consistent with our previous report that characterized the NPC mice with accelerated aging phenotype ^5^, these analyses establish that NPC1 deficiency imposes a state of markedly accelerated cellular transcriptomic aging that is partially reversible by Fingolimod. In addition, our data indicates that Fingolimod exerts a consistent, cell-type-independent aging deceleration effect at the transcriptome level.

**Figure 9.**
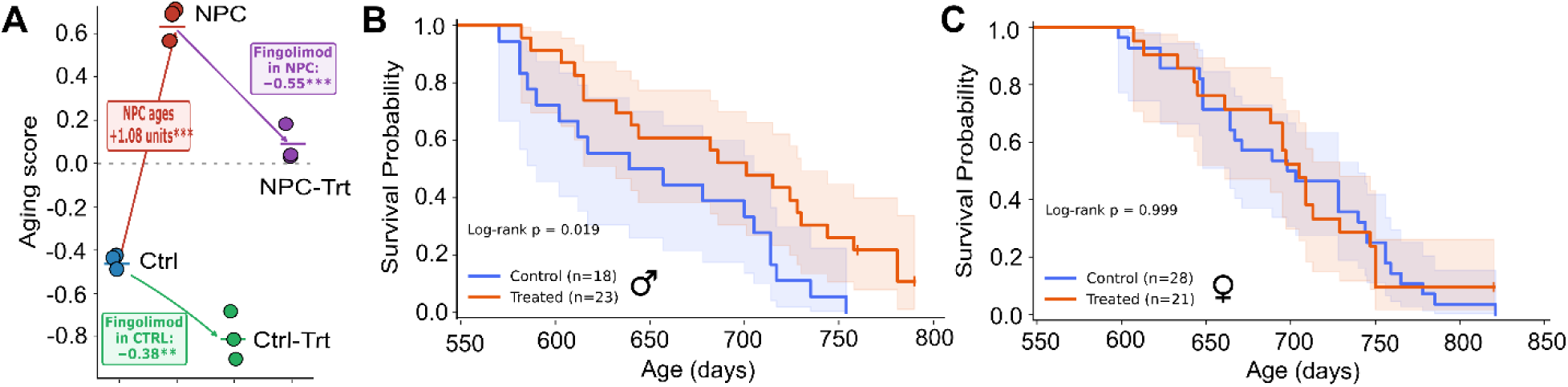
Fingolimod attenuates cellular aging and promotes organismal longevity in mice. (**A**) Transcriptome-based aging clock analysis using 100 curated aging-associated genes applied to RNA-seq datasets from health control, untreated NPC, and Fingolimod-treated NPC fibroblasts. Healthy control fibroblasts exhibited a youthful baseline aging score (−0.46 ± 0.01), whereas untreated NPC fibroblasts displayed a markedly elevated aging signature (+0.61 ± 0.03), consistent with accelerated cellular aging driven by NPC1 deficiency. Fingolimod treatment significantly reduced the NPC aging score (+0.06 ± 0.03), partially reversing the aging phenotype, although values remained higher than controls. Fingolimod also reduced the aging score in healthy fibroblasts (−0.84 ± 0.04), indicating a mild rejuvenation effect in non-diseased cells. (**B**) Late-life Fingolimod treatment extends lifespan in male but not female aged mice. Kaplan–Meier survival curves for aged BALB/cBy mice fed control chow or chow containing Fingolimod (∼2.5 mg/kg body weight/day) beginning at 16–18 months of age and followed until natural death or a humane endpoint. Survival was analyzed separately by sex, with control an Fingolimod-treated groups compared within each sex. **(A)** Male mice (control, n = 18; Fingolimod, n = 23). Fingolimod significantly extended survival, increasing median lifespan from 657 to 701 days (≈21.6 to ≈23.0 months; log-rank Mantel–Cox test, p = 0.026). **(C)** Female mice (control, n = 28; Fingolimod, n = 21). Median lifespan was unchanged by treatment (698 vs. 705 days), and the survival curves were essentially superimposable (log-rank Mantel–Cox test, p = 0.9). The x-axis begins at the treatment-initiation age (18 months); the upper axis shows the corresponding age in months. Vertical tick marks denote right-censored animals (alive at last observation); shaded bands indicate 95% confidence intervals, and the tables below each panel show the number of animals at risk. Mice that died from fighting were excluded; animals alive at the time of analysis were included as right-censored observations.

Next, to examine whether this *in vitro* anti-senescent effects of Fingolimod extend to organismal aging, we tested the effect of late-life treatment on mice lifespan. The FDA approved Fingolimod has been reported to extend lifespan when administered between 0.1 and 1mg/kg in specific disease models (e.g., neurodegeneration, lysosomal storage disorders), it has been shown to extend health- and lifespan by suppressing neuroinflammation and protecting against demyelination ^47–50^. However, there have been no reports of Fingolimod’s effects in WT mouse cohorts. Here, 16-18-month-old male and female Balb/cBy mice were fed with chow containing fingolimod corresponding at a dose of 2.5 mg/kg/day or with control chow without the drug. To further validate the Fingolimod absorption, we measured the Sphingosine 1-phosphate (S1P), sphingosine, Fingolimod and phospho-Fingolimod in the blood samples of mice fed with drug or control chow. Fingolimod acts as a functional antagonist of S1P receptors ^51^. As expected, we detected very low concentration of sphingosine in plasma, but higher amounts in the whole blood samples. S1P was about 1 µM in plasma and ∼4 µM in blood. Similarly, the Fingolimod was very low with a concentration of around 5nM and phospho-Fingolimod was below the detectable range (**Supplementary File S2**).

Overall, although the oral administration of Fingolimod in our study was within the commonly used dosing range for rodent studies ^47^, this data indicates that the actual absorption rate and blood level of Fingolimod is lower than the amount mixed in the chow. It is also possible that some of the Fingolimod might be degraded during the chow preparation. In addition, Fingolimod might be catabolized by gut microbiota, although current clinical research on the microbial breakdown of fingolimod in the human gut is limited. We should also highlight here that there have been no studies measuring the blood level of Fingolimod after oral administration in rodent studies.

Accordingly, we did not observe any side effect during the treatment and mice cohorts were maintained through the end of life or until they reached the humane end point. In males (control, n = 18; treated, n = 23), Fingolimod significantly extended survival relative to controls, increasing median lifespan from 657 to 701 days (21.6 to 23.0 months; log-rank p = 0.026) (**Fig. 9B**). In contrast, no survival benefit was observed in females (control, n = 28; treated, n = 21), where the survival curves were essentially superimposable and median lifespan was unchanged (698 vs. 705 days; log-rank p = 0.9). (**Fig. 9C**). These results indicate that late-life Fingolimod treatment confers a sex-specific extension of lifespan in male, but not female, aged mice.

Finally, to examine whether this prolongevity effect of Fingolimod is conserved across different domains of life, we analyzed lifespan in nematode worm, *Caenorhabditis elegans* (*C. elegans*) under Fingolimod treatment conditions. Fingolimod treatment led to a significant (p= 0.0002) median lifespan extension that in comparison to the control treated animals showed ∼30% longer median lifespan (Control versus treatment median lifespan = 15 and 19 days) (**Supplementary Fig. S9**). In addition, Fingolimod alleviated the age-associated phenotypes such that Fingolimod treatment significantly reduced lipofuscin accumulation, reactive oxygen species (ROS) levels, and lipid content in treated worms in comparison to the untreated controls (**Supplementary Fig. S9)**. To our knowledge, our data represents the first results that Fingolimod promotes lifespan extension and improves age-associated phenotypes in a multicellular organism.

Collectively, our multi-scale analyses demonstrate that NPC1 deficiency drives a coordinated program of transcriptomic, mitochondrial, and organismal aging, characterized by accelerated aging signatures, mitochondrial dysfunction, and shortened lifespan. Fingolimod consistently attenuates these aging-associated phenotypes across systems, partially reversing the transcriptomic aging signature, normalizing mitochondrial membrane potential and mtDNA mislocalization, and improving organismal healthspan in both invertebrate and mammalian models. Notably, the lifespan-extending effects of Fingolimod are sex-dependent in mice, with significant benefits observed in males but not females, highlighting a sex-dependent modulation of aging pathways. Together, these findings position Fingolimod as a potential geroprotector targeting conserved aging-related mechanisms linking lysosomal dysfunction, mitochondrial homeostasis, and systemic aging.

## DISCUSSION

Because of the rarity of NPC disease, progress in understanding its pathogenic mechanisms and developing effective therapies has been limited. NPC is a severe neurodegenerative disorder that affects not only the central nervous system but also peripheral organs, including the liver and spleen. Accumulating evidence indicates that neuroinflammation is a central driver of disease progression and targeting inflammation drivers such as STING has been shown to substantially ameliorate NPC pathology ^14^. In this study, we provide mechanistic insight into NPC pathogenesis by demonstrating extensive inflammation, DNA damage, and the presence of cytoplasmic mt-DNA fragments, all of which are associated with robust neuroinflammation and pronounced neuronal loss. DNA damage has been increasingly implicated in neurodegeneration, particularly in disorders of defective repair such as ataxia-telangiectasia, Cockayne syndrome, and xeroderma pigmentosum ^[3^,^56]^, and as a hallmark of brain aging more broadly ^52–55^. NPC, is known as a metabolic-trafficking disorder, and to our knowledge no systematic prior assessment of genome integrity has been reported in this disease. Our identification of widespread double-strand breaks in NPC cerebellum and patient fibroblasts therefore represents an extension of the genome-instability paradigm into monogenic lysosomal disease.

Equally novel is the chronic DNA damage tolerance phenotype we identified: induction of mutagenic translesion synthesis polymerases (*REV3L*, *POLI*, *POLK*) coupled with coordinated suppression of high-fidelity homologous recombination and base excision repair pathways. This pattern has been described in cancer cells exposed to chronic replication stress ^56,57^, but has not, to our knowledge, been documented in a lysosomal storage disorder, and it raises the intriguing possibility that long-term NPC1 deficiency fosters somatic mutational signatures contributing to progressive cellular dysfunction. Together with mitochondrial DNA puncta outside MitoTracker-defined boundaries, these observations point to a unified mechanism in which lysosomal failure generates a pool of cytosolic nucleic acid that activates the cGAS–STING pathway driving the chronic neuroinflammation ^30,58^. Of particular conceptual importance is our finding that STING protein and *TMEM173* mRNA are oppositely regulated in NPC patient derived fibroblasts. NPC1 has recently been identified not only as a lysosomal cholesterol transporter but also as a lysosomal adaptor that regulates STING degradation ^14^. Loss of functional NPC1 impairs lysosomal turnover of STING, leading to its accumulation and sustained activation. This mechanism has been reported to occur largely independently of cGAS ^14^.

Given the extensive DNA damage observed in NPC, we propose that cytoplasmic mt-DNA fragments may also contribute to STING activation, in parallel with impaired lysosomal degradation. STING activity is further modulated by intracellular cholesterol levels, STING contains two cholesterol-binding motifs that regulate its retention in the endoplasmic reticulum (ER), thereby establishing a cholesterol-dependent threshold for pathway activation ^[50^,^63]^. Depletion of ER cholesterol facilitates STING release, trafficking to the Golgi, and downstream signaling ^[50]^. In NPC1 disease, defective lysosomal cholesterol export results in cholesterol accumulation within lysosomes and concomitant depletion of ER cholesterol, a condition that primes STING activation. Thus, altered cholesterol trafficking represents a key mechanistic link between NPC1 dysfunction and pathological STING signaling ^[12^,^50]^.

Although the upstream drivers of neuroinflammation in NPC disease remain incompletely understood, our data reveal profound DNA damage and an associated cellular crisis. Nevertheless, these findings reinforce the concept that suppression of STING signaling alone is sufficient to significantly ameliorate disease progression, supporting a central role for STING in NPC pathology ^[12]^. In addition, our findings demonstrate that Fingolimod significantly increases LAMP1 protein levels, suggesting a previously unrecognized mechanism by which Fingolimod modulates STING signaling and neuroinflammation in NPC disease. To our knowledge, this is the first study showing that Fingolimod upregulates LAMP1 expression, potentially facilitating cholesterol redistribution from lysosomes and thereby limiting aberrant STING activation. Collectively, these findings support a model in which impaired lysosomal cholesterol trafficking contributes to pathological STING signaling in NPC1 disease and identify LAMP1-mediated cholesterol redistribution as a potential therapeutic mechanism for restoring lysosomal homeostasis and suppressing inflammation.

This model is consistent with recent evidence indicating that overexpression of the lysosomal membrane protein LAMP1 can partially compensate for NPC1 dysfunction by facilitating cholesterol export from late endosome/lysosome compartments toward the ER and plasma membrane ^59,60^. Elevated LAMP1 expression and consequent reduction of lysosomal cholesterol accumulation have also been proposed as underlying mechanisms contributing to the therapeutic effects of HPβ-cyclodextrin (HPβCD) in NPC disease models ^15,16,59,61^. Accordingly, targeting STING directly or modulating cholesterol transport and metabolism represents a promising therapeutic strategy for NPC. Despite these beneficial effects, Fingolimod did not reduce DNA damage or cytoplasmic DNA accumulation. These observations indicate that although targeting inflammation and lipid trafficking significantly ameliorates disease pathology, DNA damage represents an additional and independent contributor to NPC pathogenesis. The mechanistic relationship between impaired lipid metabolism, lysosomal dysfunction, mitochondrial stress, and genomic instability remains poorly understood and warrants further investigation. Therapeutic strategies that directly target DNA damage responses may therefore provide additional benefit in NPC disease.

The pronounced sex specificity of Fingolimod’s *in vivo* action is consistent with known sex differences in S1P receptor expression, sphingolipid metabolism, microglial baseline biology, and S1P-modulator efficacy in multiple sclerosis ^62–67^. A notable conceptual implication of our findings is the inverse relationship between baseline disease severity and therapeutic responsiveness: male NPC mice exhibited more severe gliosis and DNA damage than females, yet responded more robustly to Fingolimod treatment, suggesting that the magnitude of disease-associated dysregulation may influence the extent of pharmacological rescue. These observations raise the possibility that Fingolimod preferentially restores homeostasis under conditions of heightened pathological stress rather than acting through strictly sex-dependent mechanisms alone. Although the relatively small cohort size used for several analyses (n = 3 per sex per condition) limits the strength of these conclusions, the convergence of histological, behavioral, and molecular findings supports the presence of biologically meaningful sexual dimorphism and reinforces growing recognition of sex as a critical biological variable in neuroscience research ^68,69^. Notably, lifespan analyses performed in larger cohorts of aged mice (n = ∼20 per group) further supported the preferential efficacy of Fingolimod in males, consistent with the observations obtained in NPC models.

Our most far-reaching observation is that Fingolimod decelerates cellular aging and shows a conserved, positive lifespan effect: it extended median lifespan by ∼30% in *C. elegans* and by ∼7% in aged male mice. To our knowledge, these findings represent the first demonstrations of lifespan-extending and anti-aging effects for Fingolimod in multicellular organisms. This places Fingolimod alongside the small set of clinically approved drugs with cross-species lifespan-extending activity, including rapamycin ^[73]^, metformin ^[74]^, and acarbose ^[75]^. The mechanistic basis appears multimodal, engaging at least four recognized hallmarks of aging ^[76]^, including mitochondrial dysfunction (ΔΨ_m_ normalization, mtDNA confinement), cellular senescence (p21 reduction), chronic inflammation/neuroinflammation (STING level, microglia activation, astrocytes reactivation), and dysregulated lipid signaling (cholesterol mobilization and LAMP1 level). Simultaneous engagement of multiple hallmarks is the defining feature of geroprotective drugs and distinguishes Fingolimod from agents targeting individual aging axes. Importantly, Fingolimod is FDA-approved with an established safety profile, oral bioavailability, and blood–brain-barrier penetrance properties ^[77^,^78]^, that distinguish it from many candidate gerotherapeutics still in early development and provide a near-term rationale for evaluation in age-related neurological conditions and, with appropriate pharmacovigilance, in healthy aging cohorts.

In summary, our study suggests that NPC disease involves interconnected pathological features spanning lysosomal cholesterol trafficking defects, chronic innate immune activation, mitochondrial dysfunction, and accelerated cellular aging. Using patient-derived fibroblasts and Npc1-mutant mouse models, we show that NPC1 deficiency is associated with widespread DNA damage and cytosolic mitochondrial DNA accumulation, which together may contribute to sustained inflammatory signaling, including activation of the STING pathway. Importantly, we demonstrate that Fingolimod partially ameliorates key disease-associated alterations in NPC, including neuroinflammatory responses, senescence-associated transcriptional programs, mitochondrial abnormalities, and transcriptomic aging signatures, while also extending organismal lifespan across multiple model systems. However, Fingolimod does not significantly reduce the underlying DNA damage burden, indicating that genome instability is likely a parallel and persistent component of the disease process. Overall, these findings support a link between lysosomal dysfunction, innate immune activation, and aging-related phenotypes such as mitochondrial dysfunction and DNA damage in NPC and suggest that Fingolimod may act as a partial disease-modifying agent for NPC and potential geroprotector against aging by targeting interconnected hallmarks of aging.

## Materials and Methods

### Ethics statement

The animal care and experimental protocols were approved by the Institutional Animal Care and Use Committee (IACUC) of the Virginia Commonwealth University (VCU) and the University of Virginia (UVA).

### Animal Husbandry and Drug Administration for Behavioral Assays

All animal studies were conducted in accordance with VCU and UVA Institutional Animal Care and Use Committee (IACUC) guidelines. Heterozygous Npc1 mutant mice (*Npc1^tm(I1061T)Dso^*) were obtained from Jackson Laboratory (Bar Harbor, ME, USA) and bred to generate homozygous Npc1 mutants (*Npc1*^−/−^) and wild-type (WT) littermates. All mice were bred and housed under non-sterile conditions, with food and water available ad lib. The genotype was determined by polymerase chain reaction (PCR) using previously reported primer sets. Fingolimod was dissolved in ethanol and was given to mice through gavage at a dose of 2 mg/kg per day starting from age 21 days to age 105 days. The measurement of body weight was performed once a week.

### Motor Function Assessment

Neurological phenotypes in NPC1^mut^ mice were monitored daily by observing tremor activity, while body weights were recorded weekly. Mice displaying neurological symptoms received Diet Gel 76A and HydroGel for adequate hydration and nutrition. To assess balance and motor coordination, mice were trained at 4 weeks of age, and a series of established phenotypic tests were performed weekly, with final data collected at 13 weeks. The Balance Beam Test required mice to traverse 60 cm on a 2 cm-diameter wooden beam within 180 seconds without falling. The Rotarod Test assessed balance and coordination by recording the ability to maintain climbing on a rotating drum (3 cm in diameter, 15 RPM) for 360 seconds.

### Tissue Immunofluorescence

Niemann-Pick disease (NPC) mice and wild-type controls at 105 days of age were anesthetized using isoflurane delivered via a vaporizer machine. Mice were transcardially perfused with phosphate-buffered saline (PBS) at a rate of 1 mL/min for 10 minutes, followed by perfusion with 4% paraformaldehyde (PFA) in PBS for fixation. Brains were carefully dissected and post-fixed in 4% PFA for 24 hours at 4 °C. The fixed brains were then cryopreserved in 30% sucrose in PBS for 48 hours. Brains were embedded in optimal cutting temperature (OCT) compound (Tissue-Tek, VWR) and stored at −80°C for 24 hours. Serial 50μm sections were prepared using a cryostat (Leica CM1860) and mounted on probe plus microscopy slides (Fisher Scientific).

Sections were permeabilized in PBS containing 1% Triton X-100 for 30 minutes at room temperature. To block non-specific binding, sections were incubated in blocking buffer, containing PBS with 0.5% Triton X-100 and 10% goat serum for 1 hour at room temperature. Primary antibodies used included anti-GFAP (Cell Signaling; 1:300) for astrocytes, anti-IBA1 (Wako Chemicals; 1:1000) for microglia, anti-Calbindin (Abcam; 1:500), Anti-γH2AX (Cell Signaling; 1:100) and anti-cGAS (Cell Signaling; 1:100). Antibodies were diluted in blocking buffer and incubated with tissue sections overnight at 4°C. The following day, sections were washed three times with PBS and incubated with the secondary antibodies including Alexa Fluor 594, 633 and 488 (Invitrogen; 1:500) for 45 minutes at 37°C. Sections were washed three times with PBS and mounted using Vectashield mounting medium (Vector Laboratories). Slides were imaged using a Zeiss LSM 880 microscope at 20x and 40x magnification. Maximum projection images from z-stacks were obtained, ensuring no fluorescence crossover between channels. Image analysis was performed using ImageJ software.

### Cell Culture and Drug Treatment

Fibroblast cells derived from healthy control (Coriell Institute accession number AG04438) and NPC patients (accession number GM18436) were obtained from the Coriell Institute. Cells were cultured in α-minimum essential medium (α-MEM) supplemented with 15% fetal bovine serum (FBS) and 1% penicillin–streptomycin. Cells were maintained at 37°C in a humidified atmosphere containing 5% CO. Media were changed every 2–3 days, and cells were passaged at approximately 80–90% confluence using 0.05% trypsin–EDTA for detachment. For treatment, cells at 60–70% confluence were exposed to Fingolimod at a final concentration of 25nM. After 24 hours, the medium was replaced with fresh medium containing the same concentration of Fingolimod, and cells were incubated for an additional 24 hours.

### Western blotting

Cell lysates were prepared using RIPA buffer containing 1% protease inhibitor, followed by protein quantification with the BCA assay. Equal amounts of protein were resolved on SDS-PAGE and transferred to PVDF membranes. Membranes were blocked in 3% bovine serum albumin (BSA) and incubated overnight at 4°C with primary antibodies against p21, STING, LAMP1 and β-actin as the loading control. After washing, membranes were incubated with HRP-conjugated secondary antibodies for 1 hour. Protein bands were visualized using ECL and imaged with the ChemiDoc XRS+ system. Densitometry analysis was conducted with ImageJ, normalizing protein expression to β-actin.

### Cellular Immunofluorescence

Cells were fixed with 4% PFA for 15 min at room temperature, washed three times with PBS, and permeabilized with 0.2% Triton X-100 in PBS for 10 min. Following blocking with 5% BSA in PBS for 1 h, cells were incubated overnight at 4 °C with primary antibodies against γH2AX (Cell Signaling, 1:100) diluted in blocking solution. After washing, cells were incubated for 1 hour at room temperature with Alexa Fluor-conjugated secondary antibody (Invitrogen, 1:1000). Nuclei were counterstained with DAPI (1 µg/mL, Sigma-Aldrich). Samples were mounted in antifade mounting medium and imaged using Cytation 10 imaging platform (Agilent BioTek)

### Cholesterol Level Detection Using Filipin III

Unesterified cholesterol was detected using Filipin III dye (Sigma-Aldrich). Cells were fixed with 4% PFA for 15 min, washed with PBS, and incubated with Filipin III working solution (50 µg/mL in PBS) for 1 h at room temperature in the dark. After washing with PBS, stained cells were imaged using the Cytation 10 Cell Imaging Multi-Mode Reader (Agilent BioTek).

### Mitochondrial Imaging and Functional Assessment

Mitochondrial membrane potential (ΔΨm) was assessed using MitoTracker CMXRos Red (Thermo Fisher Scientific, M7512; 50 nM). Cells were washed three times with Hank’s Balanced Salt Solution (HBSS), then incubated with a staining master mix containing MitoTracker CMXRos Red (50 nM), PicoGreen (Thermo Fisher Scientific, P11496; 1:500 dilution), and Hoechst 33342 (Thermo Fisher Scientific, 62249; 5 µM) for 30 minutes at 37°C. Following staining, cells were washed three times with HBSS and allowed to equilibrate for 10 minutes at 37°C before imaging. Fluorescence intensity per cell was quantified using ImageJ (NIH).

Super-resolution live-cell imaging was performed on a Zeiss LSM 980 confocal microscope equipped with an Airyscan 2 detector. Cells were maintained at 37°C with 5% CO in a stage-top incubator throughout image acquisition. Cells were stained as described above with MitoTracker CMXRos Red (50 nM; mitochondrial membrane potential), PicoGreen (1:500 dilution; mitochondrial DNA), and Hoechst 33342 (5 µM; nuclei) prepared as a master mix. Cells were washed three times with HBSS, incubated with the dye mixture for 30 minutes at 37°C, washed three times with HBSS, and equilibrated for 10 minutes at 37°C in the incubator prior to imaging. Images were processed using Zeiss ZEN software. Statistical analyses were performed using GraphPad Prism. Data are expressed as mean ± standard error of the mean (SEM). Comparisons across groups were analyzed by Brown-Forsythe and Welch one-way analysis of variance (ANOVA), which does not assume equal variances across groups, followed by Dunnett’s T3 post hoc multiple comparisons test. A p-value of < 0.05 was considered statistically significant.

### Transcriptome-based aging clock and aging score computation

To quantify the transcriptome-wide aging phenotype of each sample, we constructed a composite transcriptomic aging score using a curated panel of 100 aging-associated genes, 64 genes whose expression increases with chronological age (aging-UP genes) and 36 genes whose expression decreases with chronological age (aging-DOWN genes), derived from the integration of four independently validated resources. First, the cross-tissue transcriptomic aging gene signature of which identified 1,497 genes whose expression changes significantly with age across multiple human tissues in a meta-analysis of 7,074 individuals, was used as the primary source for tissue-validated aging-up and aging-down genes ^70^. Second, the SenMayo senescence gene set, comprising 125 genes that define the senescence-associated secretory phenotype across species, was incorporated as the reference for SASP and pro-senescence effectors ^71^. Third, the GenAge Human Aging Gene Database (HAGR; https://genomics.senescence.info/genes), which curates genes associated with human longevity, progeroid syndromes, and age-related disease, was used to define the longevity and anti-aging gene set^72^. Fourth, the CellAge cellular senescence database (https://genomics.senescence.info/cells), which catalogs 279 genes that regulate replicative and stress-induced cellular senescence, was incorporated to capture cell-cycle arrest and senescence regulatory transcripts ^73^. Gene set membership was further annotated by pathway according to the nine Hallmarks of Aging framework^46,74^: (1) genomic instability, (2) telomere attrition, (3) epigenetic alterations, (4) loss of proteostasis, (5) disabled macroautophagy, (6) deregulated nutrient sensing, (7) mitochondrial dysfunction, (8) cellular senescence, and (9) altered intercellular communication. All gene symbols were standardized to HGNC nomenclature and verified against Ensembl release 111. The final panel comprised 100 genes after retaining only those measurably expressed across the dataset.

The transcriptomic aging score was computed on a per-sample basis as follows. For each of the 100 aging genes, log_2_(expression + 1) values were extracted from the relevant source comparison file, yielding a gene × sample expression matrix comprising all twelve samples simultaneously. Each gene was z-scored across all twelve samples to remove absolute expression differences between genes:

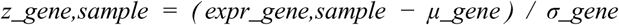

where μ_gene_and σ_gene_ are the mean and standard deviation of log_2_(expression + 1) across all twelve samples for that gene. A pseudovalue of 10^−8^ was added to σ to prevent division by zero. The per-sample aging score was then computed as:

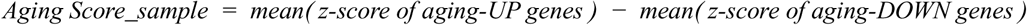

A positive score indicates a transcriptome skewed toward the aging direction (higher expression of pro-aging genes, lower expression of longevity genes); a negative score indicates a transcriptome skewed toward the youthful direction. This formulation is conceptually equivalent to the single-sample gene set enrichment (ssGSEA) approach applied to opposing gene modules, and is methodologically analogous to the aging score frameworks described in included datasets^70,75^.

Group-level mean aging scores and standard errors of the mean (SEM) were computed for all four experimental groups. Statistical comparisons between groups were performed using the two-sample Welch t-test. The percentage of NPC aging acceleration reversed by Fingolimod was computed as (score_NPC_ − score_NPC_T_) / (score_NPC_ − score_CTRL_) × 100.

### Genome-wide DSB mapping/sequencing and DSB read processing

DNA double-strand break detection was performed as described previously ^[32]^. Briefly, genomic DNA was purified by gently lysing cells in 50 mM Tris pH 8.0, 100 mM EDTA, 100 mM NaCl, 1% SDS, and 1 mg/ml proteinase K at 55°C for 3 hrs, followed by organic solvent extraction and ethanol precipitation. Precautions such as gentle pipetting with wide-open pipette tips was taken to avoid DNA shearing and introduction of artificial DNA breaks during purification. Purified genomic DNA was subjected to blunting/A-tailing reactions, Illumina P5 adapter ligation to capture broken DNA ends, and subsequent reactions as described previously. Prepared libraries were subjected to whole-genome 150-bp paired-end sequencing with the Illumina NovaSeq X Plus platforms. Paired end sequencing reads were mapped to the human genome (GRCh38/hg38) via Bowtie2 (v.2.3.4.1) aligner running in high sensitivity mode (*--very-sensitive*), as described previously. Unmapped, non-primary chromosome, and low-quality reads were filtered out. PCR duplicates were marked, and the first mate of non-duplicated pairs was extracted for continued analysis. For each identified break, the most 5’ nucleotide of the first mate defined the DNA break position. Two technical replicates for each sample showed high reproducibility of genomic coverage (Pearson’s correlation r = 0.97) and were subsequently combined for downstream data analysis. This strong correlation confirms that the break mapping procedure does not introduce significant amounts of random DNA breaks that could convert single-stranded nicks into DSBs.

### Consensus G4-forming regions

To generate G4 consensus sites publicly available BG4 ChIP-seq and CUT&Tag data with associated input controls from NHEK, K562, HaCaT, U2O2S, HEK293T and NSC were downloaded through Sequence Read Archive (**Supplementary Table 1**) and aligned to the GRCh38/hg38 genome using bowtie2. Biological duplicates were merged, low quality, not primary alignment and unmapped reads were filtered out and blacklist regions were removed. For each cell line G4 peaks were called individually by macs2 tool with default settings with each dataset controlled for the matching input data. G4 consensus sites were generated by combining G4 peaks from each cell line (union, n = 65 923) using BEDtools merged and then using BEDtools intersect only G4 sites that were present in at least five out of six cell lines were kept resulting in n = 5 572 genomic regions.

### Mouse Lifespan Analyses

BALB/cBy mice were obtained from the NIA Aged Rodent Colony. Experimental animals were housed under standard conditions and monitored daily throughout the study. For drug administration, custom chow was prepared by incorporating 12.5 mg fingolimod hydrochloride, 99.94% purity (MedChemExpress, Cat. No. HY-12005), into 1 kg of open standard diet (Research Diets, D11112201). This formulation was designed to deliver approximately 2.5 mg/kg body weight/day, based on an average body weight of 20 g per mouse and an estimated daily food intake of 4 g per mouse. Pellets were dried at approximately 70°C and gamma-irradiated at 10–20 kGy before packaging. Diets were stored at 4°C until use.

Experimental animals were assigned to receive either regular control diet or fingolimod-containing diet for the duration of the study and were followed until natural death or until they reached a moribund state requiring euthanasia. Mice that died as a result of fighting were excluded from survival analyses. Animals that were alive at the time of analysis were included as right-censored observations using the last observation date and were not counted as death events. Survival curves were generated separately for male and female mice using the Kaplan–Meier method, with control and fingolimod-treated groups compared within each sex. Median survival was calculated from the Kaplan–Meier estimates. Statistical differences between survival curves were assessed using the log-rank Mantel–Cox test, with p < 0.05 considered statistically significant. All mouse experiments were performed in accordance with institutional guidelines for the care and use of laboratory animals and were approved by the University of Virginia School of Medicine Institutional Animal Care and Use Committee.

### C. elegans lifespan and health span assays

Wild-type N2 *C. elegans* were maintained at 22°C on nematode growth medium (NGM) plates seeded with *E. coli* OP50. Age-synchronized populations were generated by bleaching gravid adults and allowing embryos to develop to the young adult stage. Young adult worms were transferred to NGM plates containing FUdR (150 µM; Sigma, cat no: F0503). Fingolimod was prepared as a stock solution in DMSO and added to heat-killed OP50 to a final concentration of 500 µM; control plates received equivalent volume of DMSO. Approximately 20–30 worms were placed on each plate, with three biological replicates per condition. Lifespan was monitored using the WormBot automated imaging system ^76^, with survival scored based on movement following automated stimulation and longitudinal image-based detection. Worms were considered dead when movement was absent across consecutive imaging sessions. Median and maximum lifespan were calculated for each condition, and statistical significance was determined using log-rank test, corrected with Benjamini–Hochberg correction.

For lipid accumulation, age-synchronized N2 worms were treated with fingolimod or vehicle beginning at the young adult stage and maintained at 22°C on heat-killed OP50-seeded NGM plates until day 14 of adulthood. Worms were collected in M9 buffer, washed three times to remove residual bacteria, and incubated with Nile Red staining solution in M9 for 60 min at room temperature in the dark. Following staining, worms were washed three times with M9, briefly destained in fresh M9, immobilized with sodium azide, and imaged immediately using a fluorescence microscope with a red fluorescence filter set. Images were acquired using identical exposure, gain, and illumination settings across all groups. At least 20 worms were imaged per condition per biological replicate. Fluorescence intensity was quantified using ImageJ/Fiji, normalized to vehicle-treated controls, and analyzed using an unpaired two-tailed t-test.

Reactive oxygen species levels were measured in day-14 adult N2 worms using H2DCFDA fluorescence imaging. Worms treated with fingolimod or vehicle from the young adult stage were collected in M9 buffer and washed three times to remove residual bacteria. Worms were incubated with H2DCFDA staining solution in M9 for 60 min at room temperature in the dark, washed three times with M9, immobilized with sodium azide, and imaged immediately using a fluorescence microscope with a FITC/GFP filter set. All images were acquired using identical exposure, gain, and illumination settings. Fluorescence intensity was quantified using ImageJ/Fiji from at least 20 worms per condition per biological replicate, normalized to vehicle-treated controls, and analyzed using an unpaired two-tailed t-test.

Finally, age-associated lipofuscin accumulation was assessed in day-14 adult N2 worms treated with fingolimod or vehicle from the young adult stage. Worms were collected in M9 buffer, washed three times to remove residual bacteria, immobilized with sodium azide, and imaged immediately by fluorescence microscopy. Lipofuscin autofluorescence was detected using a DAPI filter set under identical imaging conditions for all groups. Fluorescence intensity was quantified using ImageJ/Fiji from at least 20 worms per condition per biological replicate. Values were normalized to vehicle-treated controls, and statistical significance was determined using an unpaired two-tailed t-test.

### RNA sequencing and analysis

Global gene expressions were analyzed by RNA sequencing. Briefly, RNA was isolated from fibroblast cells with/without fingolimod using Quick-RNA Miniprep Kit (Zymo Research). RNA quality was assessed by Agilent RNA 6000 Pico Kit (Agilent Technologies). Isolated RNA samples were then sent for sequencing. mRNA was isolated from bulk RNA using poly-T oligo-attached magnetic beads for library preparation and sequenced on the NovaSeq Illumina platform. The paired-end (150bp) raw reads were then assessed for quality using standard QA protocol and aligned using Hisat2 v2.0.5 (http://daehwankimlab.github.io/hisat2/) with the reference human genome GRCh38 to find raw counts. Differential expression analysis between comparison groups was performed using the DESeq2 package in R (1.20.0). DESeq2 provides statistical routines for determining differential expression in digital gene expression data using a model based on the negative binomial distribution. The resulting P-values were adjusted using Benjamini and Hochberg’s approach for controlling the false discovery rate. Genes with an adjusted P-value<0.05 found by DESeq2 were assigned as differentially expressed.

### Functional pathway over-representation analysis

Pathway over-representation was assessed using the one-tailed hypergeometric test, equivalent to Fisher’s exact test for over-representation. Three independent tests were performed for each pathway: enrichment among all DEGs irrespective of direction; enrichment among upregulated DEGs only; and enrichment among downregulated DEGs only. No cross-pathway multiple-testing correction was applied, as enrichment results were used for directional hypothesis generation and pathway characterization rather than formal discovery claims.

Pathway gene membership was curated from three principal sources. (i) KEGG pathway database (release 105.0, accessed April 2025; https://www.kegg.jp): lysosome (hsa04142), sphingolipid metabolism (hsa00600), autophagy (hsa04140), Toll-like receptor signaling (hsa04620), TNF signaling (hsa04668), complement and coagulation cascades (hsa04610), base excision repair (hsa03410), nucleotide excision repair (hsa03420), mismatch repair (hsa03430), homologous recombination (hsa03440), and non-homologous end joining (hsa03450). (ii) Molecular Signatures Database (MSigDB) Hallmark collection (v2023.2; https://www.gsea-msigdb.org): Epithelial–Mesenchymal Transition (HALLMARK_EPITHELIAL_MESENCHYMAL_TRANSITION), cholesterol homeostasis (HALLMARK_CHOLESTEROL_HOMEOSTASIS), unfolded protein response (HALLMARK_UNFOLDED_PROTEIN_RESPONSE), inflammatory response (HALLMARK_INFLAMMATORY_RESPONSE), interferon-alpha response (HALLMARK_INTERFERON_ALPHA_RESPONSE), and DNA repair (HALLMARK_DNA_REPAIR). All human gene symbols were standardized to HGNC nomenclature (https://www.genenames.org) and cross-referenced against Ensembl release 111 (GRCh38.p14).

### LC-MS/MS Methodology for Metabolite Analyses from Mouse Blood

S1P, internal standard (d7-S1P), sphingosine, FTY720 and phospho-FTY720 quantification were performed using a tandem quadrupole mass spectrometer (Waters Xevo TQ-S micro) with a UPLC inlet (Waters Acquity h-class+) with a reverse phase C-18 column (Waters BEH C-18 1.7 µm bead size, 2.1mm x 50mm). Our chromatography protocol, which is a modification of that described by ^77^, was a binary solvent gradient with a constant flow rate of 0.4 mL/min with a column temperature of 60°C. Mobile phase A (MPA) consisted of water/methanol/formic acid (79:20:1) while Mobile phase B (MPB) was methanol/acetone/water/formic acid (68:29:2:1). Following injection (1-9 µL) on column, a run began with 50:50 MPA: MPB for 0.5 min followed by increasing MPB linearly to 100% to 3.5 min and holding at 100% MPB for another 3 min. The column is re-equilibrated to 50:50 MPA:MPB for 1.5 min. Analytes were detected using these MRM protocols: S1P (380.1>264.4, voltages: cone 18, collision 16), deuterated d7-S1P (387.2>271.4, voltages: cone 24, collision 16), sphingosine (300.3>252.3, voltages: cone 30, collision 18), FTY720 (308.1>255.2 voltages: cone 4, collision 14), and phospho-FTY720 (388.1>290.3 voltages: cone 66, collision 10) all in ESI+ mode. Peak analysis was accomplished using Waters TargetLynx software ver. 1.4. Solvents were LCMS Optima grade (Fisher Scientific, Waltham, MA). Analyte concentrations were estimated from the ratio to the internal standard (d7-S1P) signal.

### Statistical Analysis

Unless otherwise indicated, all quantitative data are shown as mean ± S.D. from at least three independent experiments (biologic replicates), all of which were conducted in triplicates or duplicates (technical replicates). For comparison between two groups, unpaired Student’s t-test with two-tailed distribution was used. For the multiple group comparisons either a one- or two-way ANOVA, as appropriate, with Tukey or Sidak post hoc and Fisher’s exact test in R software (4.4.1) and GraphPad Prism software (10.4.0). A p-value <0.05 was considered statistically significant (*p < 0.05, **p < 0.01, ***p < 0.001, ****p < 0.0001)

## Acknowledgements

We gratefully acknowledge Kevin Lynch, Ph.D. from UVA, Department of Pharmacology for his assistance with the quantification of FTY720 and S1P levels in mouse blood samples. J.N. was supported by the Eunice Kennedy Shriver National Institute of Child Health and Human Development (R00HD096117) and the VCU Parkinson’s and Movement Disorders Center Pilot Research Program.

