## SUpplementalry Figures for "NPC1 deficiency engages a lysosome–genome–immune program linked to neurodegeneration and cellular aging signatures"

**Supplementary Figures**

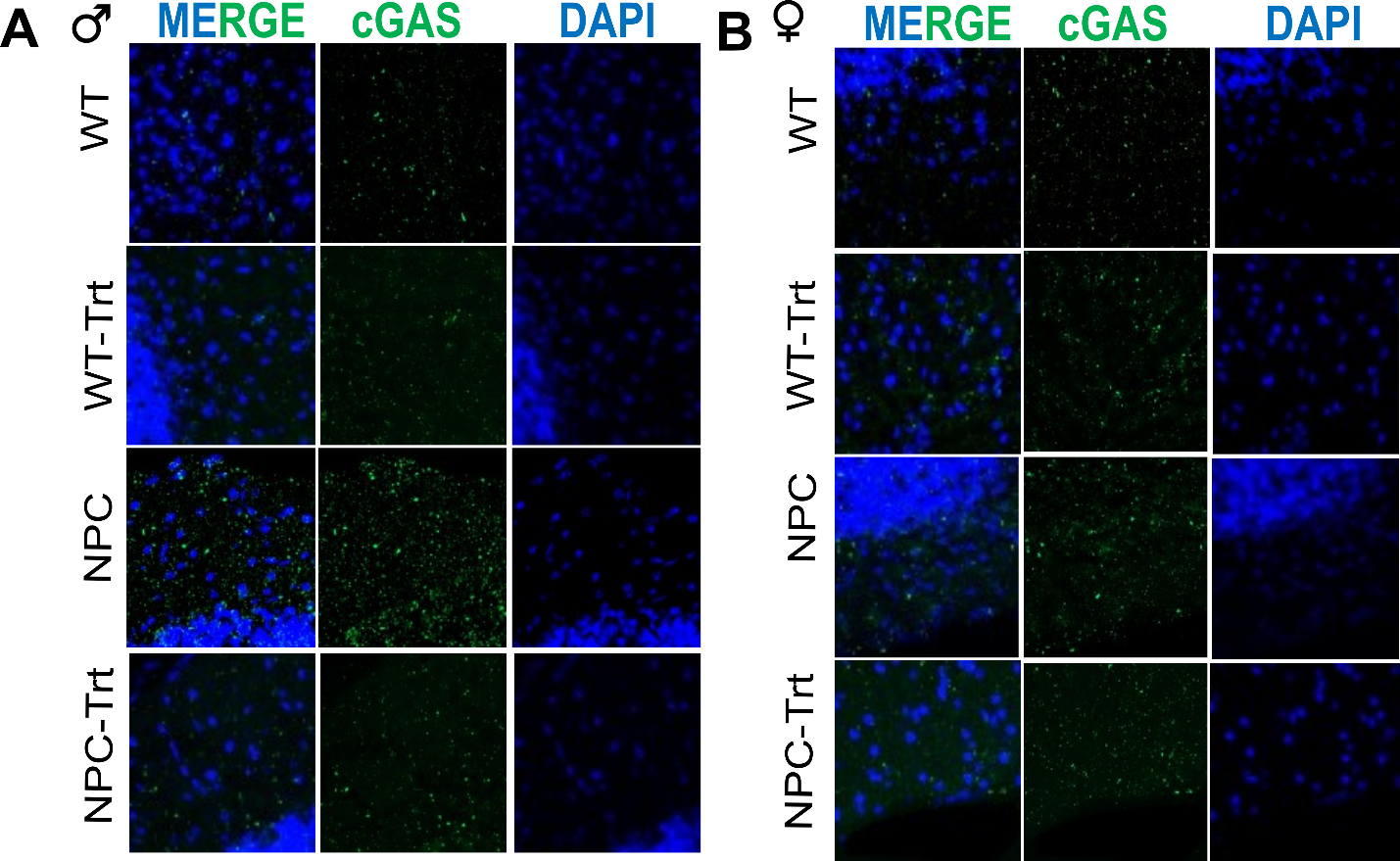

**Figure S1:** **NPC1mut mice is characterized with increased expression of cGAS.** Confocal images of molecular layer of the cerebellum from male (**A**) and female (**B**) mice for cGAS (green) and nuclei (DAPI, blue) staining. Experimental groups include wild-type (WT), WT treated with fingolimod (WT-Trt), NPC1mut (NPC), and NPC1mut treated with fingolimod (NPC-Trt).

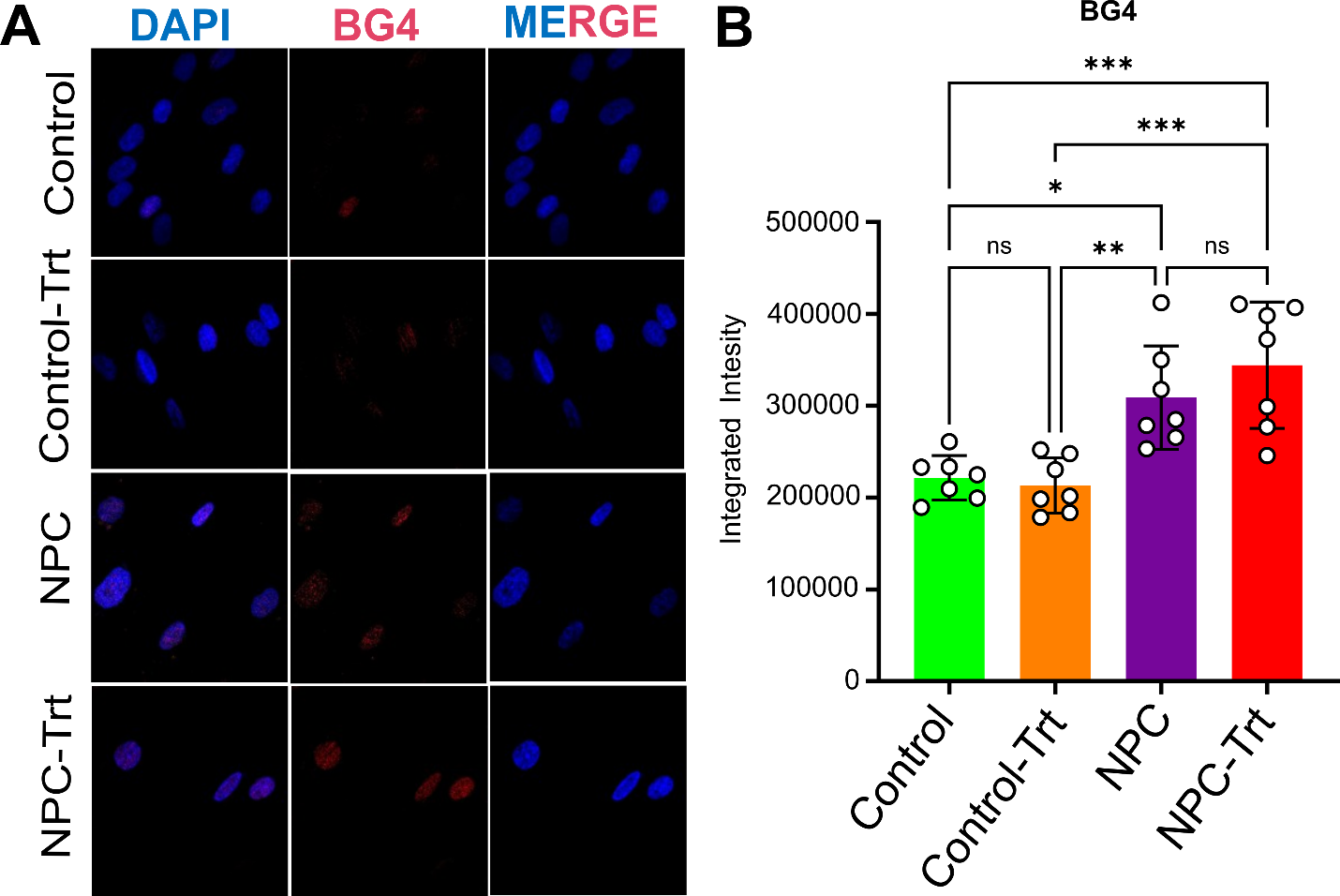

**Figure S2. NPC patient-derived fibroblast cells are quantified with elevated BG4 staining.** (**A**) G-quadruplex (G4)-specific antibody staining (red) revealed increased G4 formation in NPC cells. Experimental groups include wild-type (WT), WT treated with fingolimod (WT-Trt), NPC1mut (NPC), and NPC1mut treated with fingolimod (NPC-Trt). (**B**) Quantification shows a significant increase in G4 formation in NPC1 cells compared to control cell lines. Fingolimod did not alter the significantly alter the G4 abundance in both control and NPC cells.

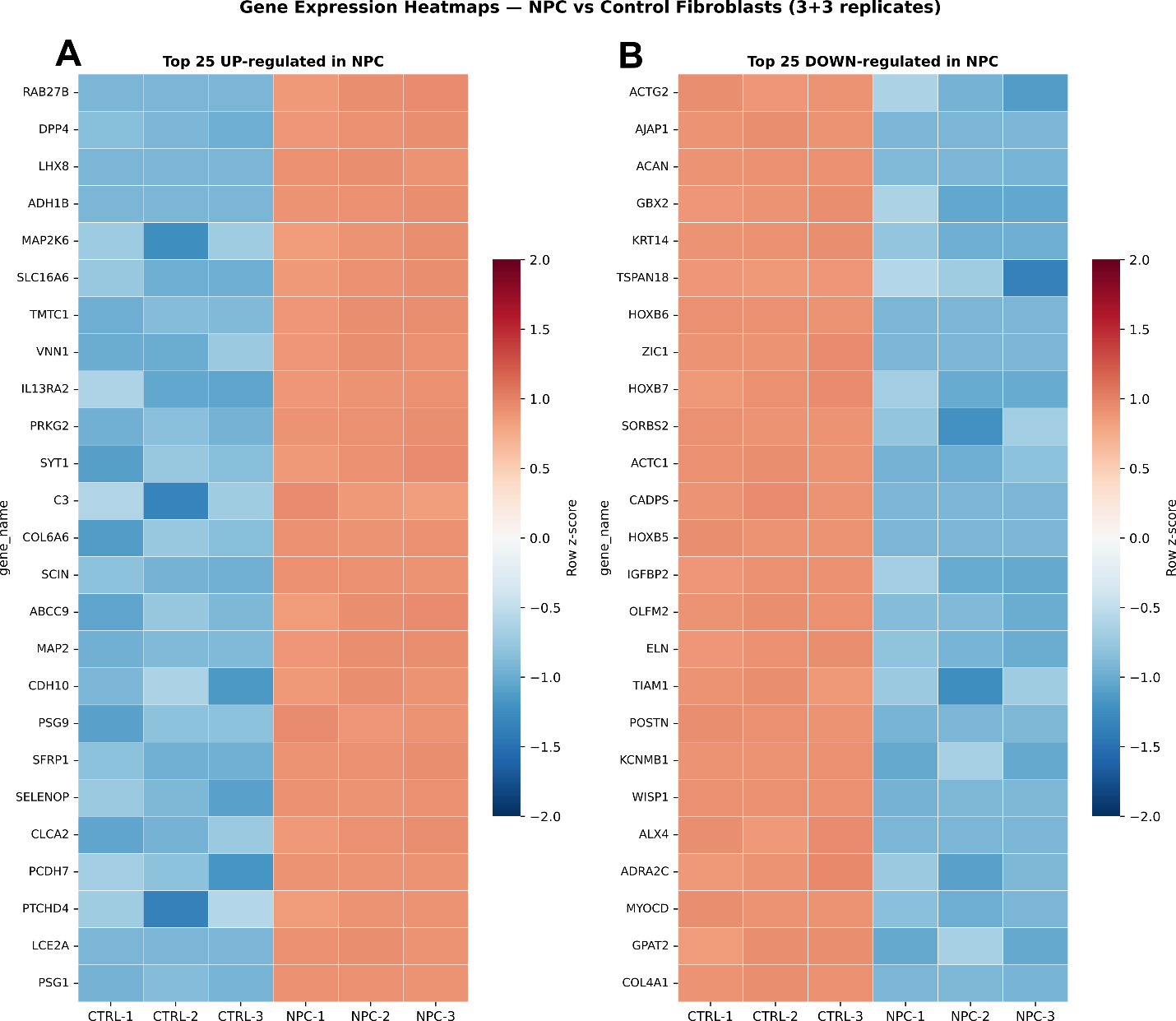

**Figure S3. Expression heatmaps of the most strongly altered genes in NPC fibroblasts.** Heatmaps showing row-normalized (z-score) log_2_-transformed expression values for the top 25 most strongly upregulated (left panel) and top 25 most strongly downregulated (right panel) genes, ranked by |log_2_FC|. Individual values for all six samples (CTRL-1, CTRL-2, CTRL-3, NPC-1, NPC-2, NPC-3) are displayed. Color scale: red indicates above-mean expression; blue indicates below-mean expression within each gene (row); scale is clamped at ±2 standard deviations. Tight clustering of replicates within each condition confirms high reproducibility of the transcriptomic profiles.

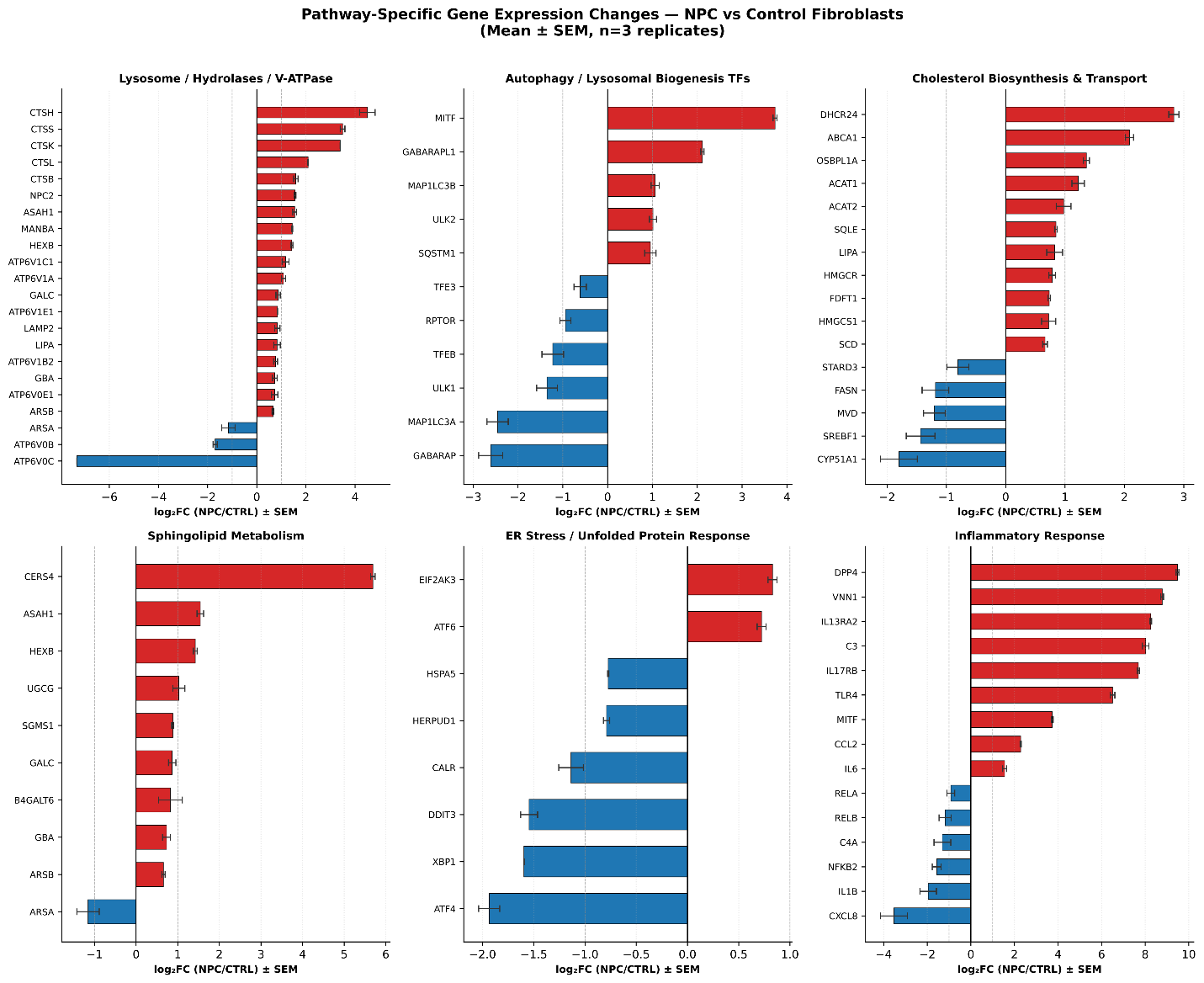

**Figure S4. Pathway-specific gene expression changes in six NPC-relevant biological modules.** Horizontal bar charts displaying log_2_FC (NPC/control) values for individual genes in six pathways. Panels: (**A**) Lysosome and V-ATPase; (**B**) Autophagy and lysosomal biogenesis transcription factors (TFEB, TFE3, MITF); (**C**) Cholesterol biosynthesis and transport; (**D**) Sphingolipid metabolism; (E) ER stress and unfolded protein response; (**F**) Inflammatory response. Red bars: upregulated in NPC; blue bars: downregulated. Grey dashed vertical lines at ±1 indicate the |log_2_FC| = 1 threshold. All genes shown are significant DEGs (adj. p < 0.05). For pathways with more than 22 members, the 11 most strongly UP and 11 most strongly DOWN genes are displayed.

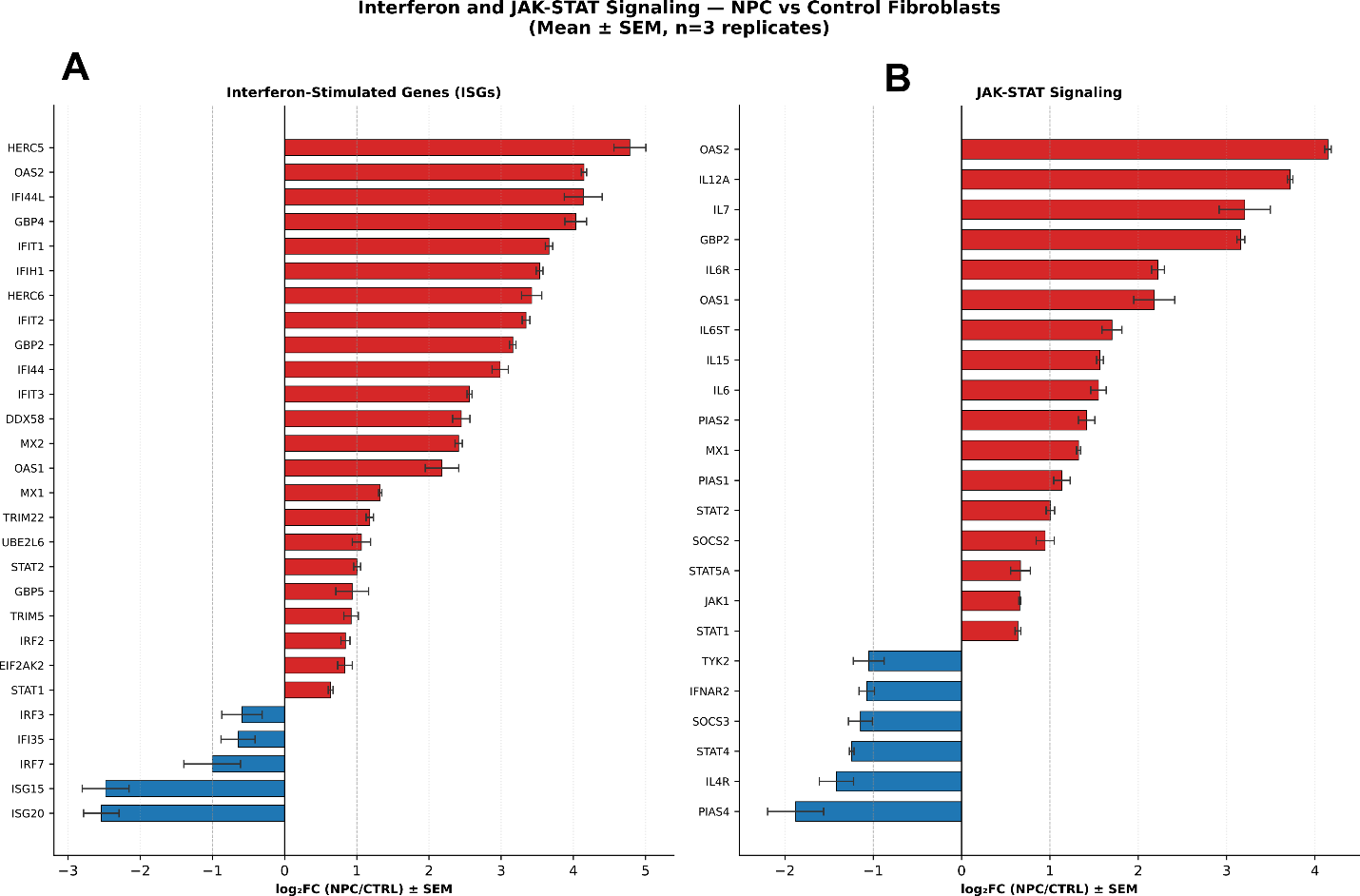

**Figure S5. Interferon-stimulated gene program and JAK-STAT signaling in NPC fibroblasts.** Bar charts of log_2_FC values for (**A**) interferon-stimulated genes (ISGs) and (B) JAK-STAT pathway members. Twenty-seven of 36 curated ISGs are significantly upregulated (hypergeometric p = 1.3 × 10^−14^), In the JAK-STAT panel, STAT2, STAT1, IL12A, IL15, and IL6R are upregulated while the feedback inhibitor SOCS3 is downregulated, implying sustained STAT signaling.

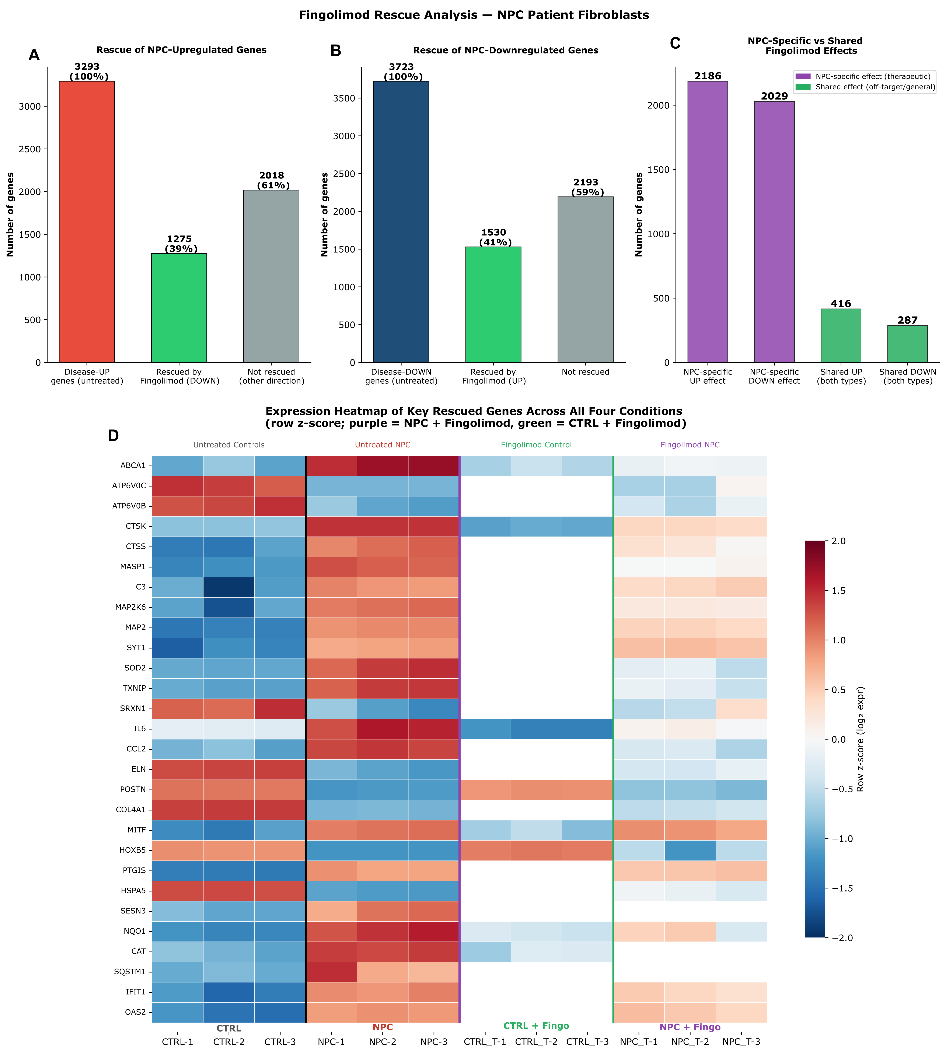

**Figure S6. Rescue gene analyses** (**A**) Of 3,293 NPC-upregulated disease genes, 1,275 (38.7%) are significantly downregulated in NPC-T vs NPC true rescue of pathologically elevated transcripts (**Top Left Panel**). Of 3,723 NPC-downregulated disease genes, 1,530 (41.1%) are significantly upregulated in NPC-T vs NPC restoration of suppressed programs (**Top Middle Panel**). 4,215 NPC-T vs NPC changes are absent in CTRL-T vs CTRL (NPC-specific), distinguishing therapeutic from off-target effects (**Top Right Panel).** (**B**) Row z-score expression heatmap across all four biological groups: CTRL (n=3), NPC (n=3), CTRL + Fingolimod (n=3), NPC + Fingolimod (n=3). Vertical lines separate the groups. Rescue is visible as genes that show high contrast between CTRL and NPC columns returning toward CTRL-like levels in the NPC + Fingolimod column. The green arrow marks the direction of ‘therapeutic normalization.

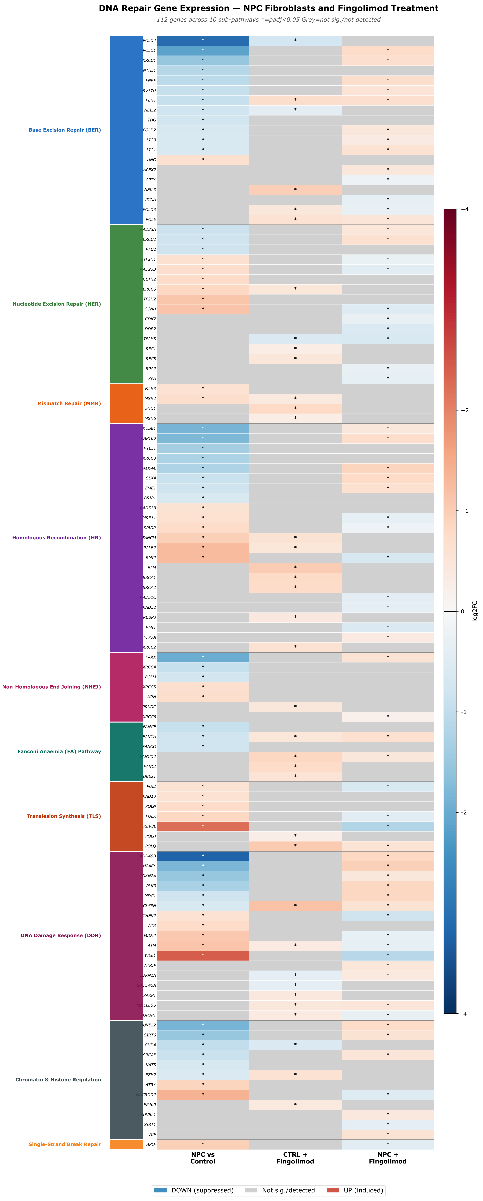

**Figure S7.** **DNA repair gene analyses.** Comprehensive heatmap of all 112 DNA repair genes detected across any comparison, grouped by sub-pathway with color-coded labels. The heatmap shows the left-right asymmetry between NPC baseline and Fingolimod responses, and the bidirectionality of the NPC+Fingo response. Within each sub-pathway group, genes are ordered by their log₂FC value in the NPC versus Control comparison. Cell color reflects the log₂ fold change on a diverging Red–Blue colourmap centered at zero (red = upregulated; blue = downregulated; scale bar shown on right; range −4 to +4. Grey cells indicate genes that were either not expressed above the detection threshold or did not reach statistical significance (BH padj ≥ 0.05) in that particular comparison and are shown for contextual completeness. An asterisk (*) within a coloured cell denotes that the gene is a significant DEG (BH padj < 0.05) in that comparison. Horizontal grey lines separate sub-pathway groups.

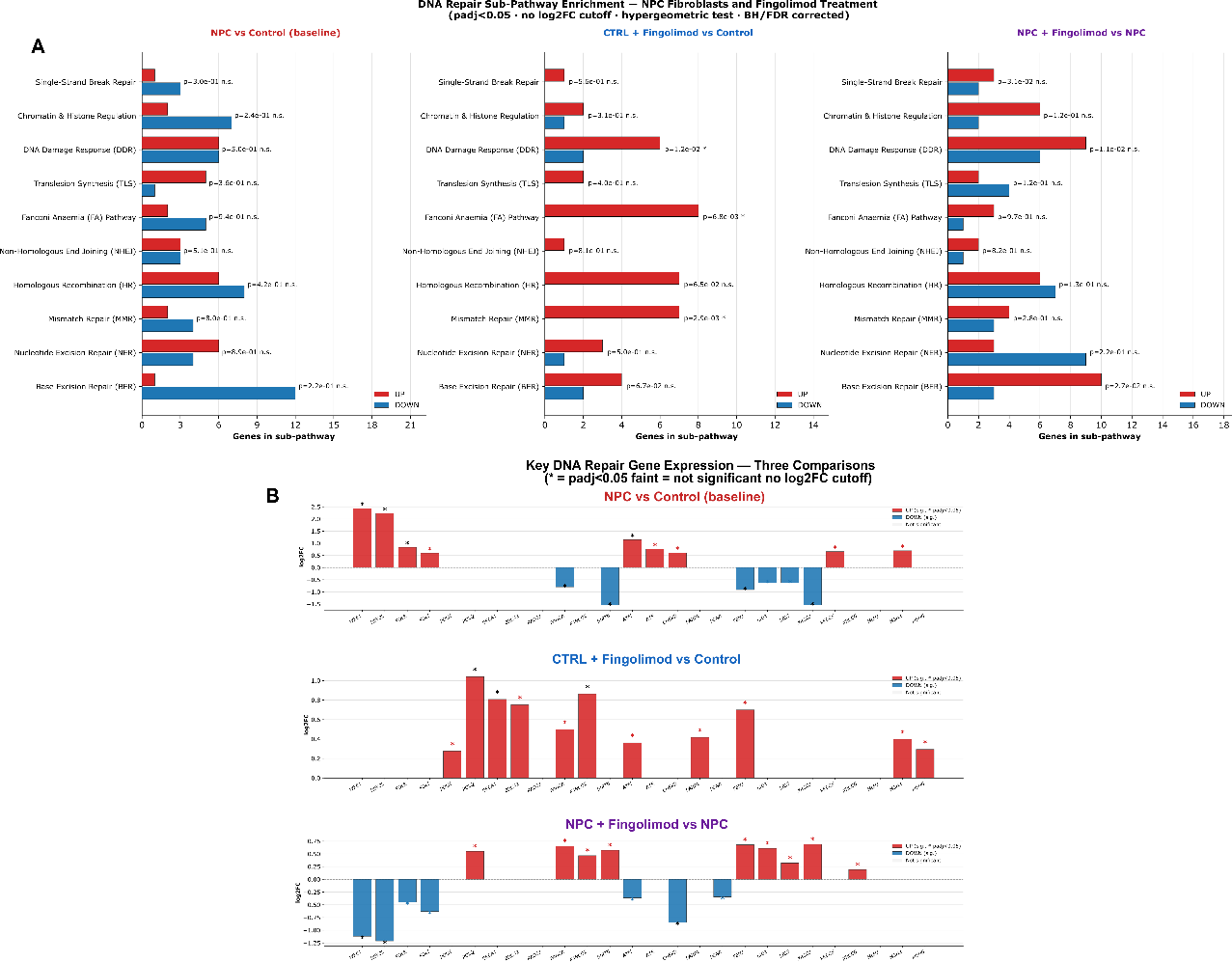

**Figure S8. DNA Repair Sub-Pathway Enrichment Analysis Across Three Pairwise Comparisons. (A)** Horizontal bar charts showing the number of significantly differentially expressed genes (DEGs; Benjamini–Hochberg adjusted p-value < 0.05, no log₂ fold change threshold applied) overlapping each of ten mechanistically defined DNA repair sub-pathways: Base Excision Repair (BER), Nucleotide Excision Repair (NER), Mismatch Repair (MMR), Homologous Recombination (HR), Non-Homologous End Joining (NHEJ), Fanconi Anaemia (FA) pathway, Translesion Synthesis (TLS), DNA Damage Response and Checkpoint signalling (DDR), Chromatin and Histone Regulation, and Single-Strand Break Repair (SSB). The DNA repair gene list comprised 200 curated, non-redundant genes assigned to sub-pathways based on established mechanistic function. Sub-pathway enrichment was assessed independently for each comparison using a one-sided hypergeometric test parameterised with a background of N = 20,000 estimated human coding genes, with all ten resulting p-values per comparison corrected simultaneously using the Benjamini–Hochberg false discovery rate procedure. Red bars represent the count of significantly upregulated (UP) genes within each sub-pathway; blue bars represent significantly downregulated (DOWN) genes. Raw hypergeometric p-values and significance annotations are displayed to the right of each bar pair (* BH-adjusted p < 0.05; ** padj < 0.01; *** padj < 0.001; n.s. = not significant after BH correction). NPC patient-derived fibroblasts versus age- and passage-matched healthy control fibroblasts (NPC vs Control). No DNA repair sub-pathway reached statistical significance after BH correction. (**A**) The absence of concentrated sub-pathway enrichment reflects the broad bidirectional dysregulation of repair genes distributed across multiple mechanisms rather than disruption of a single pathway, consistent with a NPC repair state characterized by elevated damage sensing and error-prone bypass alongside suppressed high-fidelity repair execution. CTRL + Fingolimod versus Control (CTRL+Fingo vs CTRL). Three sub-pathways were significantly UP-enriched after BH correction: Mismatch Repair (padj = 0.029), Fanconi Anaemia pathway (padj = 0.034), and DNA Damage Response/Checkpoint (padj = 0.042). All three were predominantly upregulated, consistent with a pro-proliferative, high-fidelity DNA repair programme in which Fingolimod-treated healthy cells upregulate replication-coupled repair to maintain genomic integrity during drug-induced cell cycle entry. NPC + Fingolimod versus NPC (NPC+Fingo vs NPC). No sub-pathway reached BH significance; however, the directional composition of DNA repair DEGs is qualitatively distinct from both the NPC baseline (Panel A) and the CTRL+Fingolimod response, with notable suppression of error-prone bypass genes and selective restoration of ligation and FA pathway effectors. **(B)** Three-panel bar chart showing log₂ fold change values for 26 selected DNA repair genes across treatment conditions. Genes are displayed in the same order across all three panels to facilitate direct cross-condition comparison. The y-axis represents log₂ fold change (test condition versus reference). Solid bars (red = significantly upregulated; blue = significantly downregulated; BH adjusted p < 0.05) indicate genes that are significant DEGs in that comparison. An asterisk (*) above each significant bar denotes BH padj < 0.05.

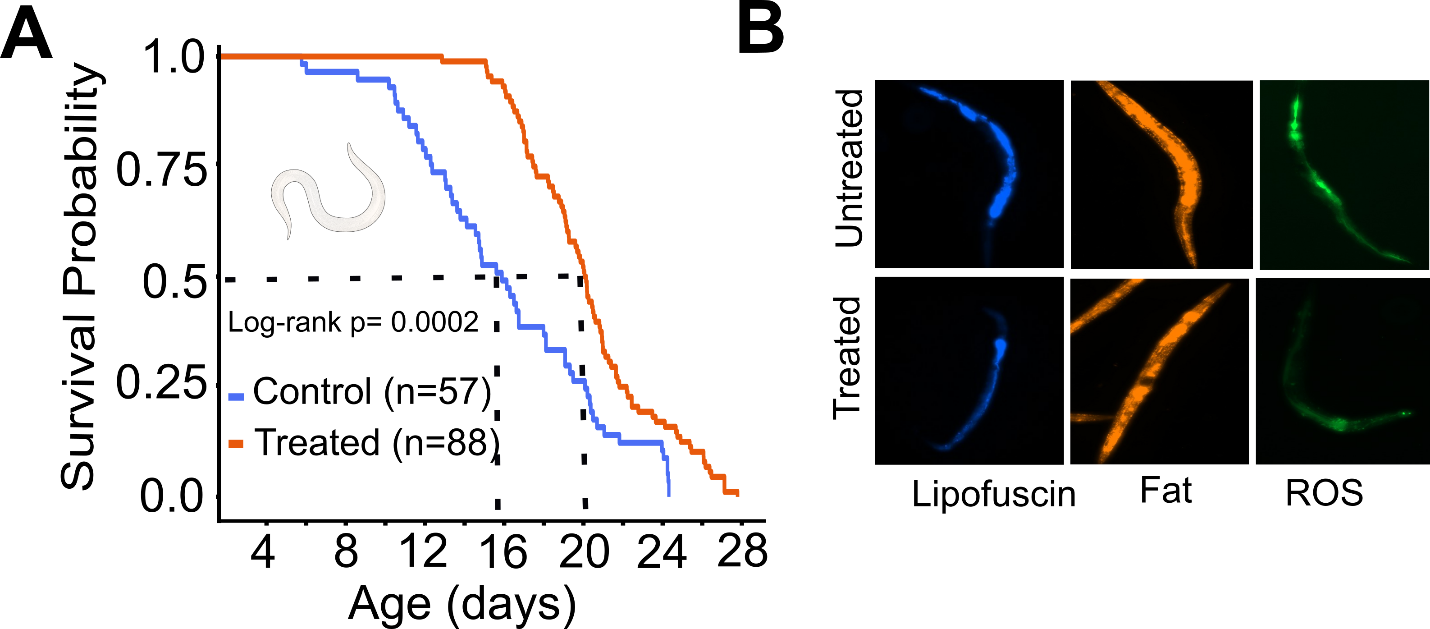

**Figure S9. *Caenorhabditis elegans* lifespan analyses.** (**A**) Fingolimod treatment significantly extends median lifespan (~30% increase) compared to controls and (**B**) improved age-associated phenotypes, including reduced lipofuscin accumulation, reactive oxygen species (ROS), and lipid content.
