## Supplementary Table for "NPC1 deficiency engages a lysosome–genome–immune program linked to neurodegeneration and cellular aging signatures"

| **Species / Cell Line** | **Dataset Type** | **Accession(s)** | **References** |
| --- | --- | --- | --- |
| *H. sapiens* / NHEK | BG4 ChIP-seq | GSE76688 | Hansel-Hertsch *et al*., Nat Genet 2016 |
| *H. sapiens* / U2OS | BG4 CUT&Tag | GSE181373 | Hui *et al*., Sci Rep 2021 |
| *H. sapiens* / K562 | BG4 ChIP-seq | GSE107690 | Mao *et al.,* Nat Struct Mol Biol 2018 |
| *H. sapiens* / HaCaT | BG4 ChIP-seq | GSE99205 | Hansel-Hertsch *et al*., Nat Protoc 2018 |
| *H. sapiens* / NSC | BG4 ChIP-seq | GSE161531 | Zyner *et al.,* Nat Comm 2022 |
| *H. sapiens* / HEK293 | BG4 CUT&Tag | GSE173103 | Lyu *et al.,* NAR 2022 |

**Supplementary Table 1**: Publicly available BG4 ChIP-seq and CUT&Tag data with associated input controls from NHEK, K562, HaCaT, U2O2, HEK293T and NSC were downloaded through Sequence Read Archive to generate G4 consensus sites.
